# Exploration and recency as the main proximate causes of probability matching: a reinforcement learning analysis

**DOI:** 10.1101/104752

**Authors:** Carolina Feher da Silva, Camila Gomes Victorino, Nestor Caticha, Marcus Vinícius Chrysóstomo Baldo

**Author notes:** Present address: Department of Psychology, University of Surrey, Guildford, Surrey, GU2 7XH, United Kingdom (Camila Gomes Victorino).

## Abstract

Research has not yet reached a consensus on why humans match probabilities instead of maximise in a probability learning task. The most influential explanation is that they search for patterns in the random sequence of outcomes. Other explanations, such as expectation matching, are plausible, but do not consider how reinforcement learning shapes people’s choices.

We aimed to quantify how human performance in a probability learning task is affected by pattern search and reinforcement learning. We collected behavioural data from 84 young adult participants who performed a probability learning task wherein the majority outcome was rewarded with 0.7 probability, and analysed the data using a reinforcement learning model that searches for patterns. Model simulations indicated that pattern search, exploration, recency (discounting early experiences), and forgetting may impair performance.

Our analysis estimated that 85% (95% HDI [76, 94]) of participants searched for patterns and believed that each trial outcome depended on one or two previous ones. The estimated impact of pattern search on performance was, however, only 6%, while those of exploration and recency were 19% and 13% respectively. This suggests that probability matching is caused by uncertainty about how outcomes are generated, which leads to pattern search, exploration, and recency.

## Introduction

In our lives, we frequently make decisions, some of which have lifelong consequences for our well-being. It is thus essential to identify the environmental and neurobiological factors that promote suboptimal decisions. Accomplishing this goal, however, can be hard. Sometimes decades of research are not enough to produce a consensus on why people often make poor decisions in certain contexts. One example is the binary probability learning task. In this task, participants are asked to choose repeatedly between two options—for instance, in each trial they are asked to predict if a ball will appear on the left or on the right of a computer screen—and if their prediction is correct, they receive a reward. In each trial, the rewarded option is determined independently and with fixed probabilities; for instance, the ball may appear on the left with 0.7 probability or on the right with 0.3 probability. Usually one option, called the majority option, has a higher probability of being rewarded than the other. A typical probability learning task consists of hundreds or thousands of trials, and as this scenario repeats itself, all participants must learn is that one option is more frequently rewarded than the other. Indeed, the optimal strategy, called maximising, is simply choosing the majority option every time. Human participants, however, rarely maximise; their behaviour is usually described as probability matching, which consists of choosing each option with approximately the same probability it is rewarded^1–3^. We would thus expect a participant performing our example task to choose left in about 70% of the trials and right in about 30% of trials, instead of optimally choosing left in all trials. Probability matching is suboptimal in this example because it leads to an expected accuracy of 30% × 30% + 70% × 70% = 58%, while maximising leads to an expected accuracy of 70%. (More generally, if the majority option is rewarded with probability 0.5 < *p <* 1, maximising leads to an expected accuracy of *p*, while probability matching leads to an expected accuracy of *p*^2^ + (1 − *p*)^2^, which is strictly less than *p*, because 0.5 < *p <* 1 implies *p*^2^ + (1 − *p*)^2^ = 1 − 2*p*(1 − *p*) < 1 − (1 − *p*) = *p*.) Since the 1950s, a huge number of studies have attempted to explain why people make suboptimal decisions in such a simple context, and many plausible causes have been proposed, but no consensus has yet been reached on how much each cause contributes to probability matching^1–3^.

Perhaps the most influential proposal is that probability matching reflects the well-known human tendency to see patterns in noise^4^: people may not realise that each outcome is randomly and independently drawn, but may believe instead that the outcome sequence follows a pattern, which they will then try to figure out^2, 5–11^. This pattern-search hypothesis is supported by much experimental evidence^5–9^. For instance, when researchers altered the outcome sequence in a probability learning task to make it look more random (by, oddly, making it less random), participants chose the majority option more frequently and performed better^6^. Moreover, participants who matched probabilities more closely in the absence of a pattern tended to achieve greater accuracy in the presence of one^9^.

It is not clear, however, how pattern search leads to probability matching. Wolford et al.^6^ claimed that “if there were a real pattern in the data, then any successful hypothesis about that pattern would result in frequency matching.” This assumes participants search for patterns by making predictions in accordance with plausible patterns. Koehler and James^2^, however, wondered why participants would employ such a strategy if they could, to advantage, maximise until a pattern was actually found. Maximising while searching for patterns, besides guaranteeing that a majority of rewards would be obtained, is also an effortless strategy^12^ that allows participants to dedicate most of their cognitive resources to pattern search^2^.

### Patterns and Markov chains

Plonsky et al.^13^ proposed an alternative explanation as to why searching for complex patterns leads to probability matching: it creates a tendency to base decisions on small samples of previous outcomes. This assumes a general model of pattern search that we will now explain in detail, since it was also adopted in our study. Let us first define a temporal pattern as a connection between past events and a future one, so that the latter can be predicted with greater accuracy whenever the former are known. Suppose, for instance, that in each trial of a task, participants are asked to predict if a target will appear on the left or on the right of a computer screen. If the target appears alternately on the left and on the right, participants who have learned this pattern can correctly predict the next location of the target whenever they know its previous location.

An event may be more or less predictable from previous events depending on the probability that links their occurrences. For instance, if the probability is 1 that the target will appear on one side in the next trial given that it was on the other side in the previous trial, the target will always alternate between sides. If this probability is greater than 0.5 but less than 1, the target will generally alternate between sides but may also appear more than once on the same side sequentially.

In general, the probability that each event will occur may be conditional on the occurrence of the *L* ≥ 0 previous events. Formally, this sequence of events constitutes a Markov chain of order *L*. In a typical probability learning task, for instance, the outcome probabilities do not depend on any previous outcomes (*L* = 0). In an alternating sequence, each outcome depends on the previous one (*L* = 1). As outcomes depend on an increasing number of past ones, more complex patterns are generated. It has been shown that participants can implicitly learn to exploit outcome dependencies at least as remote as three trials^14, 15^.

In explicit pattern learning tasks, it is believed that relevant information about past events is stored in working memory to allow prediction of the next event, while previously learned relationships between events are stored in long-term memory. To understand how predictive events are selected to enter working memory, a number of highly complex “Gating” models (e.g.^16–18^) were proposed. They assume that working memory elements are maintained or updated according to reinforcement learning rules. We will, however, simply assume that working memory stores the previous *k* outcomes, where *k* ≥ 0 depends on the perceived pattern complexity, and that participants try to learn the optimal action after each possible history of *k* outcomes. For instance, if working memory stores just the previous outcome (*k* = 1) and the outcome sequence generally alternates between left and right (*L* = 1), participants will eventually learn that left is the optimal prediction after right and right is the optimal prediction after left. In general, participants must store at least the *L* previous outcomes in working memory to learn the pattern in a Markov chain of order *L*, i.e., it is necessary that *k ≥ L*.

### Complex pattern search relies on small samples

Based on this general model of pattern search, Plonsky et al.^13^ proposed two specific models, the CAB-*k* and CAT models. The CAB-*k* model is the simplest one: In each trial, a simulated CAB-*k* agent considers the previous *k* outcomes and selects the action with the highest average payoff in the past, taking into account only those trials that followed the same *k* outcomes. In the example of the alternating pattern, a CAB-*k* agent with *k* = 1 will eventually learn to predict left after right (and vice versa), because predicting left had the highest average payoff in past trials that followed right (and vice versa).

In probability learning tasks, the CAB-*k* model with large *k* predicts probability matching^13^. This is because a large *k* generates long histories, which tend to occur more rarely than short ones (e.g., in a sequence of binary digits, 111 is more rare than 11). Thus, a CAB-*k* agent will base each decision only on the small number of trials that followed the rare past occurrences of the current history. More generally, making decisions based on only a small number of trials generates a bias toward probability matching. If, for example, participants were to always choose the most frequent outcome of the previous three trials and choosing left is rewarded with 0.7 probability, participants would choose left with 0.784 probability^13^. Indeed, perfect probability matching is achieved when an agent adopts a strategy known as “win-stay, lose-shift,” which consists of repeating a choice in the next trial if it resulted in a win or switching to the other option if it resulted in a loss. “Win-stay, lose-shift” may be used by participants with low working memory capacity^9^. It results in probability matching because in each trial the agent bases its decisions only on the previous outcome and simply predicts that trial’s outcome; thus, its choices and trial outcomes have the same probability distribution.

Plonsky et al.^13^ proposed that human participants search for complex patterns and make decisions based on a small number of trials. To support this proposal, they demonstrated that the CAT model, a variation of the CAB-*k* model, can reproduce a novel effect they detected in experimental data from a repeated binary choice task, “the wavy recency effect;” in particular, they demonstrated that pattern search can generate a “nonmonotonic recency effect” that is part of the wavy recency effect. They designed a task wherein selecting one of the options, the “action option,” resulted in a gain with 0.9 probability and in a loss with 0.1 probability, and selecting the other option always resulted in a zero payoff. They observed that following a loss, the frequency with which participants chose the action option actually increased above the mean for several trials, then decreased below the mean. They reproduced this effect using the CAT model with *k* = 14. With this large *k*, the negative effect of a rare loss on a CAT agent’s choice only occurs after the history of 14 outcomes that preceded the loss recurs.

However, the large *k* values proposed by Plonsky et al.^13^ to explain probability matching and the wavy recency effect in their behavioural data are inconsistent with the estimated storage capacity of the human working memory, which is of about four elements^19^. Plonsky et al.^13^ argued that their estimates are plausible because humans can learn long patterns. For instance, humans can learn the pattern 001010001100 of length 12^9^. Such a feat, however, does not imply that *k* ≥ 12; as will be demonstrated in Section “Pattern learning by MPL agents,” an agent can perfectly predict this pattern’s next digit given the previous five, which merely implies *k* ≥ 5. Similarly, in another study, researchers found evidence that in an *implicit* sequence learning task the participants’ actions were influenced by events at least five trials back^20^, but this does not imply that in an *explicit* pattern learning task participants can store more than five previous results in working memory. In general, studies of implicit pattern learning measure only the reaction time to the target’s appearance rather than the prediction accuracy, and it is usually not possible to determine if participants learned the complete pattern or extracted a simpler pattern that could still predict sequence elements with above-chance accuracy. Moreover, even if participants can store more results than the estimated capacity of working memory—by storing short sequences of results as “chunks,” for instance—the resulting learning problem may be intractable. The number of histories an agent must learn about increases exponentially with *k*, and this creates a critical computational problem known as the “curse of dimensionality”^17^. The value *k* = 14 generates 2^14^ = 16384 distinct histories of past outcomes for participants to learn about. If each history is equally likely to occur, learning the pattern would only be feasible if participants had tens of thousands of trials to learn from. In the cited study^13^, they only had a hundred.

### Expectation matching

Subsequently, Plonsky and Erev^21^ proposed the CATIE model, which introduces additional decision making mechanisms such as inertia and explains the same findings with a lower working memory usage. Moreover, both probability matching and the wavy recency effect can be explained by another proposed mechanism, known as expectation matching^2^. According to this proposal, probability matching arises when participants use intuitive expectations about outcome frequencies to guide their choices^2, 22, 23^. Participants intuitively understand that if, for example, outcome A occurs with 0.7 probability and outcome B with 0.3 probability, in a sequence of 10 trials outcome A will occur in about 7 trials and outcome B in about 3. Instead of using this understanding to devise a good choice strategy, participants use it directly as a choice heuristics to avoid expending any more mental energy on the problem; that is, they predict A in about 7 of 10 trials and B in about 3. There is compelling evidence that expectation matching arises intuitively to most participants, while maximising requires deliberation to be recognised as superior; e.g., when undergraduate students were asked which strategy, among a number of provided alternatives, they would choose in a probability learning task, most of them chose probability matching^22, 24^.

Expectation matching can also explain the wavy recency effect. In the task devised by Plonsky et al.^13^, losses occurred with 0.1 probability. If losses were to occur at regular intervals, the next loss would be expected to occur 10 trials after the previous loss, and indeed participants were most likely to select the action option a few trials after a loss and least likely about 10 or more trials after a loss. It is also well known that people have misconceptions about sequences of random independent events, and many believe that when one event occurs successively in a binary sequence, it will increase the probability that the other event will occur—the gambler’s fallacy^25, 26^. It is thus possible that, soon after a loss occurred, participants did not expect another to occur so soon and thought it safe to choose the action option, which caused the initial positive effect on choice frequency; as time went on, though, they might have believed a loss was about to recur and become more and more afraid of choosing the action option, which caused the delayed negative effect on choice frequency.

Most evidence for expectation matching, however, comes from experiments that employed tasks without trial-by-trial reinforcement and whose instructions described the process of outcome generation^2^. Participants would, for instance, be asked to guess all at once a colour sequence generated by rolling ten times a ten-sided die with seven green faces and three red faces^27^. In a probability learning task, however, participants do not know how outcomes are generated; they have to figure that out. More importantly, the probability learning task is a reinforcement learning task. Again and again, participants select an action and receive immediate feedback about their choices. When they make a correct choice, they are rewarded with money; otherwise, they fail to win money or, depending on the task, they lose money. Indeed, prediction accuracy improves with longer training and larger monetary rewards^28^ or when participants are both rewarded for their correct choices and punished by their incorrect ones, instead of only one or the other^29^. In reinforcement learning tasks, as responses are reinforced, they tend to become more habitual^30^ and thus less affected by conscious choice heuristics such as expectation matching.

### Reinforcement learning

A better explanation for probability matching in probability learning tasks may thus be one that takes into account how reinforcement learning shapes people’s choices. Already in the 1950s, probability learning was tentatively explained by a number of stochastic learning models, with updating rules based on reinforcement, which under some conditions predicted asymptotic probability matching (e.g.,^31, 32^).

More recently, reinforcement learning models based on modern reinforcement learning theory^33^, such as Q-Learning^34^, SARSA^35^, EVL^36^, PVL^37^, and PVL2^38^, have been used to describe how humans learn in similar tasks, such as the Iowa, Soochow, and Bechara Gambling Tasks^36–39^ and others (e.g.^30, 40^). Reinforcement learning models that incorporate representations of opponent behaviour have successfully explained probability matching in competitive choice tasks^41^. These models do not only describe many behavioural findings accurately but are also biologically realistic in that the signals they predict correspond closely to the responses emitted by the dopamine neurons of the midbrain (see^42–45^ for reviews).

Reinforcement learning models^36–38^ assume that agents compute the expected utility of each option, not their probabilities. They are thus incapable of explicitly matching probabilities and cannot explain why participants would consciously or unconsciously try to do so. The term “probability matching,” however, does not imply that participants are trying to match probabilities as a *strategy,* only that their average *behaviour* matches them approximately. As previously discussed, probability matching is achieved when an agent with no knowledge of the outcome probabilities adopts the “win-stay, lose-shift” strategy or searches for very complex patterns. In this work, therefore, we will focus not on why people match probabilities in a probability learning task, but more broadly on why they fail to perform optimally.

### Exploration, fictive learning, recency, and forgetting

Reinforcement learning models suggest many mechanisms that may contribute to a suboptimal performance in probability learning tasks, such as exploration. For a reinforcement learning agent to maximise its expected reward, it must choose the actions that produce the most reward. But to do so, it must first discover what actions produce the most reward. If the agent can only learn from what it has experienced, it can only discover the best actions by exploring the entire array of actions and trying those it has not tried before. It follows, then, that to find the optimal actions, the agent must *not* choose the actions that have so far produced the most reward. A dilemma is thus created: on one hand, if the agent only exploits the actions that have so far produced the most reward, it may never learn the optimal actions; on the other hand, if it keeps exploring actions, it may never maximise its expected reward. To find the optimal strategy, then, an agent must explore actions at first but progressively favour those that have produced the most reward^33^. In artificial intelligence planning, exploration is usually achieved by giving agents a propensity for acting at random. This is also how exploration is commonly implemented by cognitive models of decision making, even though it is recognised that people may not always explore at random.

Moreover, animals are not limited to learning from what they have experienced; they can also learn from what they *might* have experienced^46^. Reinforcement learning models that only learn from what they have experienced are of limited utility in research, and it is often desirable to add to such models “fictive” or “counterfactual” learning signals—the ability to learn from observed, but not experienced situations. Fictive learning can speed up learning and make models more accurate at describing biological learning. Fictive learning signals predict changes in human behaviour and correlate with neuroimaging signals in brain regions involved in valuation and choice and with dopamine concentration in the striatum^47–55^. In particular, in a probability learning task, when participants make their choices, they learn both the payoff they got and the payoff they would have gotten if they had chosen the other option. Through fictive learning, they can eliminate the need to explore: they can discover the optimal action while exploiting the action that has been so far the most rewarding.

Human learning, however, may include both fictive learning and exploration. Even though fictive learning supersedes exploration in a probability learning task, exploration is a core feature of cognition at various levels since cognition’s evolutionary origins^56^. Exploratory behaviour may be triggered, perhaps unconsciously, by uncertainty about the environment, even in situations it cannot uncover more rewarding actions. In a probability learning task, even after participants have detected the majority option, they may still believe they can learn more about how outcomes are generated and thus engage in exploration, choosing the minority option and decreasing their performance. This might happen if, for instance, participants believe that there exists a strategy that will allow them to perfectly predict the outcome sequence. As long as they have not achieved perfect prediction, they might keep trying to learn more and explore instead of exploit. And indeed, when participants were frequently told they would not be able to predict all the outcomes, their performance improved^28^. The same was observed when the instructions emphasised simply predicting a single trial over predicting an entire sequence of trials^57^. Exploration may thus be a reason why participants do not maximise.

The belief that perfect prediction is possible may also lead to the belief that the environment is non-stationary, i.e., that the Markov transition matrix that generates the outcome sequence is not constant^3^. In reinforcement learning, agents adapt to a non-stationary environment by implementing recency, a strategy in which behaviour is more influenced by recent experiences than by early ones. Recency is beneficial in a non-stationary environment because early information may no longer be relevant for late decisions^33^. In a probability learning task, payoff probabilities are constant, and early information is relevant for all later decisions, but participants may come to suspect otherwise as they try to predict outcomes and often fail.

Another mechanism that impairs performance is forgetting, or learning decay. An agent’s knowledge regarding each action’s expected utility may decay with time, which in a stationary environment worsens performance. This is distinct from recency, because recency gives more weight to new information relatively to old information, but forgetting just destroys old information. Forgetting can interact with pattern search to slow down learning in the short term and impair performance in the long term. An agent that does not search for patterns needs to learn only the utility of each option. In every trial, it may forget some past knowledge, but it also acquire new knowledge from observing which option has just been rewarded. An agent that searches for patterns, however, must store information about each possible history of past outcomes. In a trial, it will only acquire new information about one of those histories, the one that has just occurred; meanwhile, knowledge about all the other histories will decay. In particular, if the agent believes that each outcome depends on many past ones, it must learn the optimal prediction after many long histories. As long histories occur more rarely than short ones on average, knowledge about them will decay more often than increase, and the agent will have to constantly relearn what it has forgotten. It may thus never learn to maximise.

### Objectives

There are thus many plausible mechanisms for probability matching, and it is possible that human performance is affected by more than one. It is still unknown to what extent each contributes to behaviour. In this study, our primary aim was to quantify the effects of pattern search, forgetting, exploration, and recency on human performance in a probability learning task.

Our secondary aim was to estimate *k*, a measure of working memory usage in pattern search, which determines how complex are the patterns people search for. This is important because, as discussed above, searching for complex patterns impairs performance by creating a tendency to make decisions based on few past observations^13^ and by interacting with forgetting. To our knowledge, only Plonsky et al.^13^ have attempted to estimate working memory usage in a reinforcement learning task, but when they used models in which pattern search was the main cause of suboptimal choices, they predicted large *k* values that lie beyond working memory capacity and generate extremely hard learning problems.

We collected behavioural data from 84 young adult participants who performed a probability learning task wherein the majority option was rewarded with 0.7 probability. We then analysed the data using a reinforcement learning model that searches for patterns in Markov chains, the Markov pattern search (MPL) model. We first compared the MPL model to the PVL model, a reinforcement learning model previously shown to perform better than many other models at describing the behaviour of healthy and clinical participants in the Iowa and Soochow Gambling Tasks^37, 38^, and to the learning WSLS model^58^, based on the “win-stay, lose-shift” strategy. The MPL model generalises the PVL model by adding recency and pattern search to it (the PVL model already includes recency and exploration). It allowed us to estimate how many participants searched for patterns, how many previous outcomes they stored in working memory, and what was the impact of pattern search, exploration, recency, and forgetting on their performance. We also analysed our experimental data set for the presence of the nonmonotonic recency effect^13^, as it has been considered an evidence of complex pattern search, and tested whether the MPL could reproduce the observed results.

## Methods

Eighty-four young adult human participants performed 300 trials of a probability learning task wherein the majority option’s probability was 0.7. Three learning models were then fitted to the data: the PVL model, which was previously proposed and validated^37, 38^, the WSLS model^39, 58^, and the MPL model, which is proposed here and generalises the PVL model by adding forgetting and pattern search. The three models were compared for their predictive accuracy using cross-validation. The MPL model was selected and simulated both to check if it can reproduce several aspects of the participants’ behaviour and to estimate how pattern search, exploration, forgetting, and recency influence a participant’s decisions in a probability learning task.

### Participants

Seventy-two undergraduate dental students at the School of Dentistry of the University of São Paulo performed the task described below for course credit. They were told the amount of credit they would receive would be proportional to their score in the task, but scores were transformed so that all students received nearly the same amount of credit. Twelve additional participants aged 22-26 were recruited at the University of São Paulo via poster advertisement and performed the same task described below, except there was no break between blocks and participants were rewarded with money. Overall, our sample consisted of 84 young adult participants.

All participants were healthy and showed no signs of neurological or psychiatric disease. All reported normal or corrected-to-normal colour vision. Exclusion criteria were: (1) use of psychoactive drugs, (2) neurological or psychiatric disorders, (3) incomplete primary school, and (4) not finishing the experiment. No participants who finished the experiment were excluded.

All experimental protocols were approved by the Ethics Committee of the Institute of Biomedical Sciences at the University of São Paulo. The experiment was conducted in accordance with the Commitee’s directives for conducting research with human participants. Written informed consent was obtained from each participant.

### Behavioural task

Participants performed 300 trials of a probability learning task. In each trial, two identical grey squares were presented on a white background and participants were asked to predict if a black ball would appear inside the left or right square (Fig. 1). They pressed A to predict that the ball would appear on the left and L to predict that it would appear on the right. Immediately afterwards, the ball would appear inside one of the squares along with a feedback message, which was “You won 1 point/5 cents” if the prediction was correct and “You won nothing” otherwise. The message remained on the screen for 500 ms, ending the trial.

**Figure 1.**
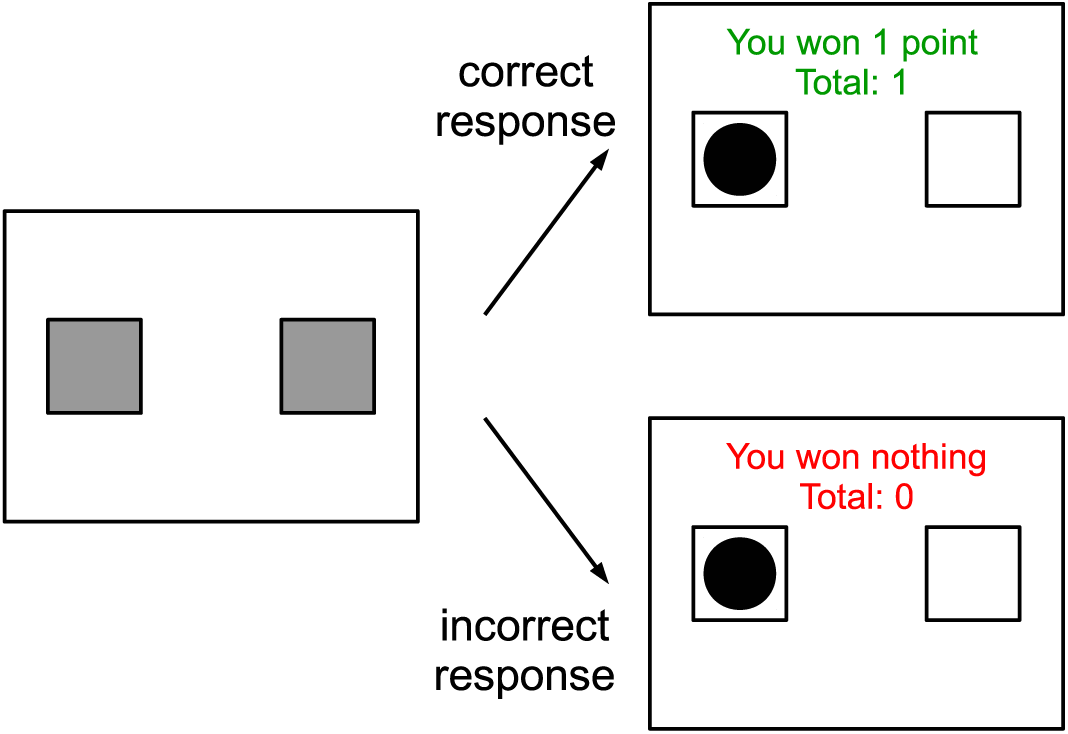
Events in a trial.

Trials were divided into 5 blocks of 60 trials with a break between them. The probabilities that the ball would appear on the right or on the left were fixed and independent of previous trials; they were 0.7 and 0.3 respectively for half of the participants and 0.3 and 0.7 for the other half. Before the task started, the experimenter explained the instructions and the participants practised them in a three-trial block. The participants did not receive any information about the structure of outcome sequences in advance.

#### Notation

The following notation will be used below: *N* is the number of participants (84) or simulated agents; for each trial *t*, 1 ≤ *t* ≤ 300, the *i*th participant’s prediction is *y_i_*(*t*) and the trial outcome is *x_i_*(*t*), where 0 and 1 are the possible outcomes; *x_i_* and *y_i_* are binary vectors containing all outcomes and predictions respectively for the *i*th participant. The majority outcome is 1, i.e., Pr[*x*(*t*) = 1] = 0.7 and Pr[*x_i_*(*t*) = 0] = 0.3, thus 1 corresponded to the left square for half of the participants and to the right square for the other half.

#### Analysis

To measure how likely participants were to choose the majority option and thus determine if they adopted a probability matching or maximising strategy, we calculated their mean response, which is equal to the frequency of choice of the majority option, since the majority option is 1 and the minority option is 0.

It has been claimed that in probability learning tasks many participants use a “win-stay, lose-shift” strategy^9, 39^. Strict “win-stay, lose-shift” implies that in each trial *t >* 1 the agent’s prediction *y*(*t*) is equal to the outcome of the previous trial *x*(*t* − 1). To check if our participants employed this strategy, we measured the proportion of responses made in accordance with “win-stay, lose-shift” by calculating the cross-correlation *c*(*x, y*) between the sequences *y* and *x* in the last 100 trials of the task, given by:

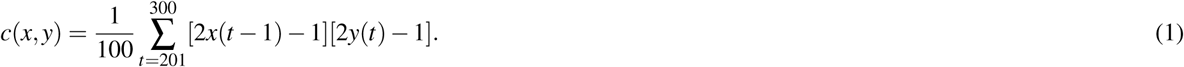

The cross-correlation is the average of [2*x*(*t* − 1) − 1][2*y*(*t*) − 1], which is equal to 1 if *x*(*t* − 1) = *y*(*t*) and to −1 if *x*(*t* − 1) ≠ *y*(*t*). If *c*(*x, y*) = 1, all predictions are the same as the previous outcome, which identifies strict “win-stay, lose-shift,” and if *c*(*x, y*) = −1, all predictions are the opposite of the previous outcome, which identifies strict “win-shift, lose-stay.”

We also investigated the “nonmonotonic recency effect”^13^. The task originally employed to investigate it had an option that resulted in a rare loss, and the task employed here did not, but it had option 1, which resulted in a gain with 0.7 probability and in a relative loss, corresponding to the missed opportunity of obtaining a gain, with 0.3 probability. It was thus possible we would also observe the nonmonotonic recency effect in our data set, and we tested for this possibility.

We adapted to our study the analysis method proposed by Plonsky et al.^13^: for every participant, trials were grouped according to the number of trials since the most recent *x* = 0 (rare) outcome; that is, for trial *t*, if trial *t − n, n >* 0, was the most recent trial with a 0 outcome, the number of trials since the most recent 0 outcome was *n*. For each participant *i* and each number of trials *n*, 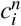 was the number of trials in the respective trial group and 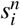 the sum of all predictions *y* in those trials, or how many times participants chose 1. The distribution of 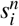 was Binomial 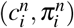, where 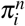 was the probability of *y* = 1. For each *n*, the parameters 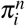 were given a beta distribution with parameters *a_n_* and *b_n_*, which were in turn given weak Half-Cauchy(0, 10^2^) prior distributions. This statistical model was coded in the Stan modelling language^59, 60^ and fitted to the data using the PyStan interface^61^ to obtain samples from the posterior distribution of model parameters from 4 chains of 30,000 iterations (warmup 15,000). Convergence was indicated by 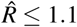 for all parameters, and at least 100 independent samples per sequence were obtained^62^. For each *n*, the participants’ mean response *a_n_/*(*a_n_* + *b_n_*) was obtained, as well as its 95% high posterior density interval (HDI).

Pattern search involving the *k* previous outcomes generates a nonmonotonic recency effect in the data, such that the mean response after a 0 outcome in trial *t* should increase in trials *t* + 1 to *t* + *k*, decrease in trial *t* + *k* + 1, then slowly increase^13^. Alternatively, a similar effect may be caused by expectation matching. If participants believed that 0 outcomes occurred regularly in the outcome sequence, they would have expected a 0 to occur every 3 to 4 trials (with 1/3 ≈ 0.33 to 1/4 = 0.25 probability), because the probability of 0 was 0.3. According to this hypothesis, the mean response should decrease three or four trials after the last 0 outcome.

We ran this analysis both in the first 100 trials of the task and in the last 100, because the nonmonotonic recency effect was first detected in a 100-trial task^13^ and, if it was caused by expectation matching, it might exist only in the beginning of the task, since over time reinforced responses are expected to become more habitual and less affected by cognitive biases such as expectation matching.

### Statistical models

Three learning models were fitted to the behavioural data: the PVL model^37, 38^, the WSLS model^39, 58, 63^, and the MPL model. The MPL model generalises the PVL model by the addition of recency and pattern search.

#### PVL model

The PVL and PVL2 reinforcement learning models have been previously evaluated for their ability to describe the behaviour of healthy and clinical participants in the Iowa and Soochow Gambling Tasks^37, 38^. They were compared to and found to perform better than many other reinforcement learning models and a baseline Bernoulli model, which assumed that participants made independent choices with constant probabilities. In this work, we adapted the PVL model to the probability learning task and used it as a baseline for comparison with the MPL model, which generalises the PVL model and is described next. The difference between the PVL and PVL2 models is not relevant for our study, since it concerns how participants attribute utility to different amounts of gain and loss. Thus we will refer only to the PVL model. The adapted PVL model combines a simple utility function with the decay-reinforcement rule^37, 38, 64^ and a softmax action selection rule^33^.

In every trial *t* of a probability learning task, a simulated PVL agent predicts the next element of a binary sequence *x*(*t*). The agent’s prediction *y*(*t*) is a function of *E*_0_(*t*) and *E*_1_(*t*), the expected utilities of options 0 and 1. Initially, *E_j_*(1) = 0 for all outcomes *j ∈* {0, 1}. The probability *p*_1_(*t*) that the agent will choose option 1 in trial *t* is given by the Boltzmann distribution:

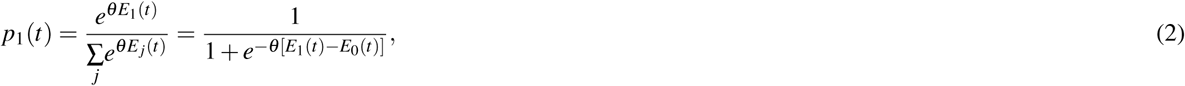

where *θ* ≥ 0 is an exploration-exploitation parameter that models the agent’s propensity to choose the option with the highest expected utility. When *θ* = 0, the agent is equally likely to choose either option (it explores). Conversely, as *θ* → ∞ the agent is more and more likely to choose the option with the highest expected utility (it exploits). The probability of a PVL agent predicting 1 in trial *t* is, as Equation 2 indicates, a logistic function of *E*_1_(*t*) − *E*_0_(*t*) with steepness *θ*. If the difference *E*_1_(*t*) − *E*_0_(*t*) is 0, i.e., both options have the same expected utility, the agent is equally likely to choose 1 or 0 (*p*_1_(*t*) = 0.5); if it is positive, the agent is more likely to choose 1 than 0, and if it is negative, the agent is more likely to choose 0 than 1.

After the agent makes its prediction and observes the trial outcome *x*(*t*), it attributes a utility *u_j_*(*t*) to each option *j*, given by:

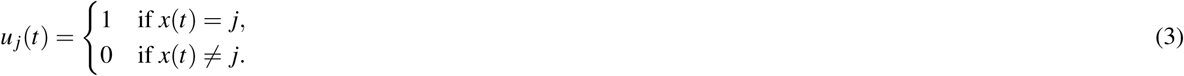

All expected utilities are then updated as follows:

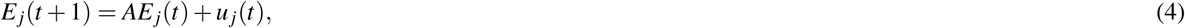

where 0 ≤ *A* ≤ 1 is a learning decay parameter, which combines both forgetting and recency.

In comparison with previous PVL and PVL2 model definitions^37, 38^, we have made two changes to adapt this model to our task. The PVL and PVL2 models were previously used to study the Iowa and Soochow Gambling Tasks, in which participants may experience different gains and losses for their choices and only learn the outcome of the choice they actually made. In our task, conversely, participants gained a fixed reward for all their correct predictions and never lost rewards; moreover, since outcomes were mutually exclusive, participants learned both the outcome of the choice they made and the outcome of the choice they could have made. To account for these differences between the tasks, we omitted the PVL features that deal with different gains and losses from the utility function and, following Schulze et al.^41^, added fictive learning to the decay-reinforcement rule.

#### MPL model

The Markov pattern learning (MPL) model uses reinforcement learning mechanisms to learn patterns in Markov chains. For a demonstration of how this model works, see Section “Pattern learning by MPL agents” below.

The MPL model includes the same two parameters per participant as the PVL model, *A* and *θ*, which measure forgetting and exploration respectively, and adds two more parameters, *k* and *ρ*, which measure working memory usage in pattern search and recency respectively. Indeed, the MPL model with *k* = 0 (no pattern search) and *ρ* = 1 (no recency) is identical to the PVL model. It is also equivalent to the CAB-*k* model^13^ with *A* = 1 (no forgetting), *ρ* = 1 (no recency), and *θ* → ∞ (no exploration).

In a probability learning task, each trial outcome *x*(*t*) is independently generated with fixed probabilities for every *t* and thus the outcome sequence constitutes a Bernoulli process. The MPL model, however, assumes that each outcome depends on the *k* previous outcomes, i.e., the outcome sequence is a Markov chain of order *k*. For every possible history (subsequence) *η* of *k* previous outcomes, the MPL agent estimates the utilities of predicting 0 or 1 in the next trial. For *k* = 2, for instance, the agent estimates the utilities of predicting 0 or 1 depending on whether the previous two outcomes were *η* = 00, *η* = 01, *η* = 10, or *η* = 11. In other words, it learns by reinforcement the Markov transition matrix of order *k* assumed to have generated the outcome sequence.

The MPL model’s utility function is identical to that of the PVL model (see above). Then, for every trial *t* and history *η* of *k* outcomes, the MPL agent computes option *j*’s expected utility 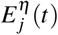. The expected utility of each option depends on the history of *k* outcomes that preceded it, and for every trial the MPL agent computes 2^*k*^ expected utilities for each option, since there are 2^*k*^ distinct histories of *k* outcomes. For instance, if *k* = 1, in each trial and for each option the agent computes two expected utilities, one if the previous outcome was 1 and another if it was 0. Initially, 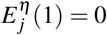 for all options *j* and histories *η*.

The agent’s next choice *y*(*t*) is a function of 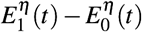. where *η* is the observed history, i.e., the *k* previous outcomes that actually occurred: {*x*(*t − k*), *x*(*t − k* + 1),…, *x*(*t* − 1)}. The probability *p*_1_(*t*) that the agent will choose option 1 in trial *t* is given by the Boltzmann distribution:

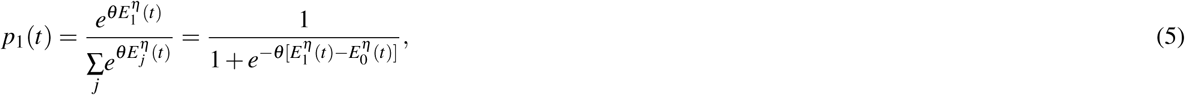

where *θ* ≥ 0 is the exploration-exploitation parameter.

After the agent makes its choice, all expected utilities referring to all histories and outcomes are updated as follows:

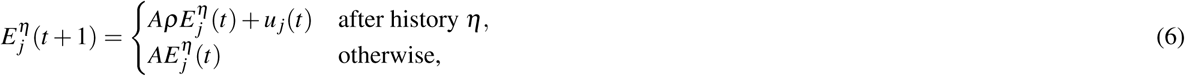

where 0 ≤ *A* ≤ 1 is a decay (forgetting) parameter and 0 ≤ *ρ* ≤ 1 is a recency parameter.

The *A* parameter is thus applied to all expected utilities associated with all possible histories, while the *ρ* parameter is only applied to the expected utilities associated with the history that actually occurred and the agent received new information about. Thus, the agent’s knowledge spontaneously decays at rate *A*, while early experiences are overridden by the most recent information at rate *ρ*. A low *ρ* value is adaptive when the environment is non-stationary and early experiences become irrelevant to future decisions. Both *A* and *ρ* cause learning decay, but if *k >* 0, they have a distinct effect on performance, as demonstrated in Section “Predicted effect of pattern search, exploration, and recency on learning speed and mean response.” If *k* = 0, there is only one possible history (the null history), which precedes every trial, and therefore all expected utilities decay at rate 0 ≤ *Aρ* ≤ 1, in which case the MPL model is identical to the PVL model with learning decay *Aρ*.

Forgetting (*A <* 1) combined with searching for complex patterns (large *k*) decreases performance. This is because the value 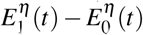 only increases after history *η*, if 1 was the outcome. Whenever history *η* does *not* occur, on the other hand, 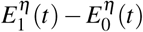 decays at rate *A*, which decreases the probability of choosing the maximising option after history *η*. As *k* increases, longer histories are generated, which occur more rarely on average, providing many opportunities for 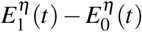 to decrease and few for it to increase.

Table 1 demonstrates how an MPL agents learns a repeating pattern for two different parameter sets.

**Table 1.** MPL agents learn an alternating pattern. MPL agents learn a sequence of outcomes *x* generated by alternating deterministically between 0 and 1. The agent’s parameters are given in the first row. The *p*_1_ column gives the probability that the agent will respond 1 (it will respond 0 with probability 1 − *p*_1_). From trial *t* = 4 on, both agents have already learned the pattern. Henceforth, the agent with optimal parameters (left) always makes correct predictions, but the agent with suboptimal parameters (right) may not always do so.

| MPL $k = 1, A = 1, \rho = 1, \theta \rightarrow \infty$ | | | | | | | MPL $k = 1, A = 0.9, \rho = 0.9, \theta = 0.3$ | | | | | | |
| --- | --- | --- | --- | --- | --- | --- | --- | --- | --- | --- | --- | --- | --- |
| $t$ | $\eta = 0$ | | $\eta = 1$ | | $p_1$ | $x$ | $t$ | $\eta = 0$ | | $\eta = 1$ | | $p_1$ | $x$ |
| | $E_0$ | $E_1$ | $E_0$ | $E_1$ | | | | $E_0$ | $E_1$ | $E_0$ | $E_1$ | | |
| 1 | 0 | 0 | 0 | 0 | 0.5 | 0 | 1 | 0 | 0 | 0 | 0 | 0.5 | 0 |
| 2 | 0 | 0 | 0 | 0 | 0.5 | 1 | 2 | 0 | 0 | 0 | 0 | 0.5 | 1 |
| 3 | 0 | 1 | 0 | 0 | 0.5 | 0 | 3 | 0 | 1 | 0 | 0 | 0.5 | 0 |
| 4 | 0 | 1 | 1 | 0 | 1 | 1 | 4 | 0 | 0.9 | 1 | 0 | 0.57 | 1 |
| 5 | 0 | 2 | 1 | 0 | 0 | 0 | 5 | 0 | 1.73 | 0.9 | 0 | 0.43 | 0 |
| 6 | 0 | 2 | 2 | 0 | 1 | 1 | 6 | 0 | 1.56 | 1.73 | 0 | 0.61 | 1 |
| 7 | 0 | 3 | 2 | 0 | 0 | 0 | 7 | 0 | 2.26 | 1.56 | 0 | 0.39 | 0 |
| 8 | 0 | 3 | 3 | 0 | 1 | 1 | 8 | 0 | 2.03 | 2.26 | 0 | 0.65 | 1 |

#### MPL parameter recovery

Since in this study we made conclusions about the participants’ strategies based on MPL model parameters, we first tested how much information about the parameters it was possible to extract from sequences of 300 predictions and outcomes generated by 10,000 simulated MPL agents. The agents had random parameters: *k* was drawn from a uniform distribution in the set {0, 1, 2, 3, 4, 5}, *A* and *ρ* were drawn from a uniform distribution in [0,1], and *θ* was drawn from a uniform distribution in [0,5]. For each agent, the resulting data were analyzed with a Bayesian model where the prior distributions of MPL parameters were the same distributions used to obtain them. Since the optimal way of updating a probability distribution with data is given by Bayes’s theorem, this Bayesian model was optimal for this analysis—it would recover the parameters as well as possible. The model was coded in the Stan modelling language^59, 60^ and fitted to the simulated data using the PyStan interface^61^ to obtain 4 chains of 3500 iterations (warmup: 1000) from the posterior distribution of model parameters. Convergence was indicated by 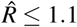 for all parameters. We then calculated the Kullback–Leibler divergence between the prior and posterior distributions of MPL parameters, which measures how much information about the parameters could be recovered from the data.

#### WSLS model

The PVL and MPL models can themselves generate “win-stay, lose-shift” behaviour. This strategy implies a tendency to choose the previous outcome, which is created by setting *k* = 0 (no pattern search) and *Aρ* = 0 (only the most recent outcome influences decisions), since with these parameter values, the expected utility of the previous outcome is always 1 and that of the other option is always 0. If *θ* → ∞ (no exploration), the agent will employ a “win-stay, lose-shift” strategy strictly; otherwise, it will employ it probabilistically.

Nevertheless, since several previous studies suggest that many participants use a “win-stay, lose-shift” strategy^9, 39^, we also compared the PVL and MPL models to a model directly inspired by this strategy, namely the learning “win-stay, lose-shift” (WSLS) model^58^. (We also tested the simplest WSLS model^58^, but it performed worse than the learning WSLS model and is not discussed further.)

In every trial *t*, the learning WSLS model assigns a probability *p_w_*(*t*) that the agent will stay (i.e., *y*(*t−*1) = *y*(*t*)) after a win (i.e., *y*(*t−*1) = *x*(*t−*1)) and a probability *p_l_*(*t*) that the agent will shift (i.e., *y*(*t−*1) ≠ *y*(*t*)) after a loss (i.e., *y*(*t*−1)≠*x*(*t*−1)). Thus, the probability *p*_1_(*t*) that the participant will chose 1 in trial *t* is given by

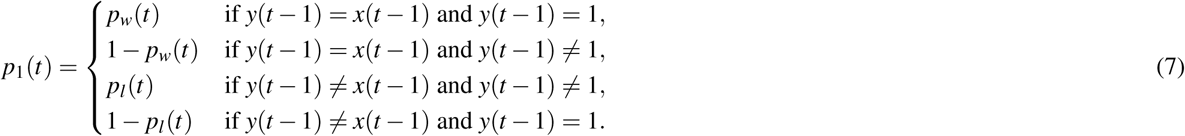

The parameters of the model are the initial probability of staying after a win *p_w_*(1), the initial probability of shifting after a loss *p_l_*(1), and two learning rates *θ_w_* and *θ_l_* for learning *p_w_*(*t*) and *p_l_*(*t*) respectively. All parameters have values in the [0, 1] interval. Learning occurs in each trial according to the following equations:

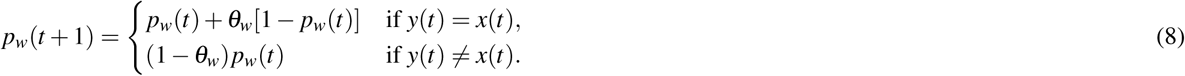

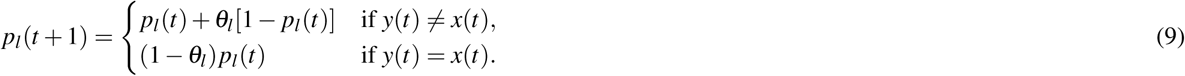

### Bayesian hierarchical models

The results of the MPL parameter recovery analysis, discussed in the Results section, show that it was not possible to recover precise information about a single simulated agent’s parameters from experimental data when the *θ* parameter had a small value, which seemed to be the case for the human participants. For this reason, the PVL, MPL, and WSLS models were fitted to each participant as part of larger Bayesian hierarchical (multilevel) models, which included the PVL, MPL, or WSLS distributions of each participant’s predictions as well as a population distribution of model parameters. This allowed us to use data from all participants to improve individual parameter estimates, to estimate the distribution of parameters across participants, and to make inferences about the behaviour of additional participants performing the probability learning task. Most of this study’s conclusions were based on such inferences. Moreover, a hierarchical model can have more parameters per participant and avoid overfitting, because the population distribution creates a dependence among parameter values for different participants so that they are not free to assume any value^62^. This was important for the present study, since the MPL and WSLS models are more complex than the PVL model, having four parameters per participant instead of two.

For each participant *i*, the PVL model has two parameters (*A_i_, θ_i_*). The vectors (logit(*A_i_*), log(*θ_i_*)) were given a multivariate Student’s *t* distribution with mean *µ*, covariance matrix Σ, and four degrees of freedom (*ν* = 4). This transformation of the parameters *A* and *θ* was used because the original values are constrained to the interval [0,1] and the transformed ones are not, which the *t* distribution requires. The *t* distribution with four degrees of freedom was used instead of the normal distribution for robustness^62^.

Based on preliminary simulations, the model’s hyperparameters were given weakly informative (regularising) prior distributions. Each component of *µ* was given a normal prior distribution with mean 0 and variance 10^4^, and Σ was decomposed into a diagonal matrix *τ*, whose diagonal components were given a half-normal prior distribution with mean 0 and variance 1, and a correlation matrix Ω, which was given an LKJ prior^65^ with shape *ν* = 1^59^.

In short, the hierarchical PVL model fitted to the experimental data was:

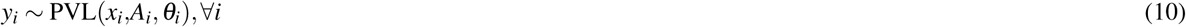

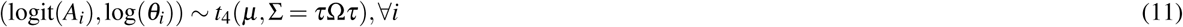

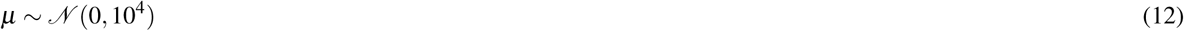

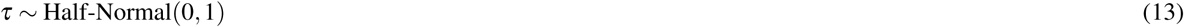

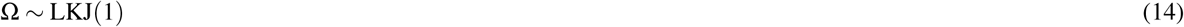

For each participant *i*, the MPL model has four parameters (*k_i_, A_i_, ρ_i_, θ_i_*). The vectors (logit(*A_i_*), logit(*ρ_i_*), log(*θ_i_*)) were given a multivariate Student’s *t* distribution with mean *µ*, covariance matrix Σ, and four degrees of freedom (*ν* = 4). The parameter *k* was constrained to the range 0–5, which is consistent with current estimates of human working memory capacity^19^. An MPL agent with working memory *k* is not limited to learning patterns of length *k*: it can also learn much longer patterns. An agent with *k* = 5, for instance, can learn the pattern 001010001100 of length 12; see Section “Pattern learning by MPL agents” for a demonstration. The parameter *k* was given a categorical distribution with Pr(*k_i_* = *k*) = *q_k_* for 0 ≤ *k* ≤ 5. In practice, the MPL model was fitted at the individual level as a mixture, with *k* as the latent variable.

The model’s hyperparameters were given weakly informative prior distributions. Each component of *µ* was given a normal prior distribution with mean 0 and variance 10^4^, and Σ was decomposed into a diagonal matrix *τ*, whose diagonal components were given a half-normal prior distribution with mean 0 and variance 1, and a correlation matrix Ω, which was given an LKJ prior with shape *ν* = 1. The hyperparameters *q_k_* for 0 ≤ *k* ≤ 5 were given a joint Dirichlet prior distribution with concentration parameter *α* = (0.001, 0.001, 0.001, 0.001, 0.001, 0.001), implying that the prior probabilities that *k* = 0, 1*,…,* 5 were 1/6.

In this hierarchical model, parameters were estimated for each participant taking into account not only which values fitted that participant’s results best, but also which values were the most frequent in the population. If, for instance, *k_i_* = 5 fitted the *i*th participant’s results best, but all the other participants had *k* ≤ 3, the estimated value of *k_i_* might be adjusted to, say, *k_i_* = 3.

In summary, the hierarchical MPL model is:

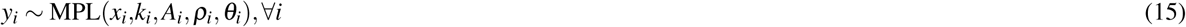

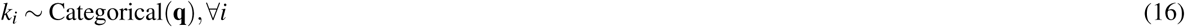

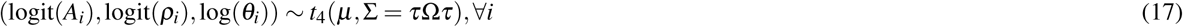

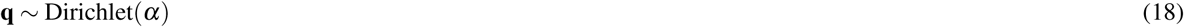

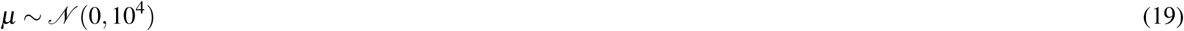

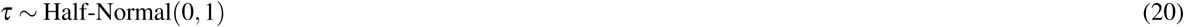

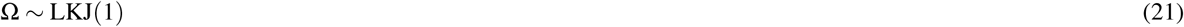

The model is also shown in Fig. 2.

**Figure 2.**
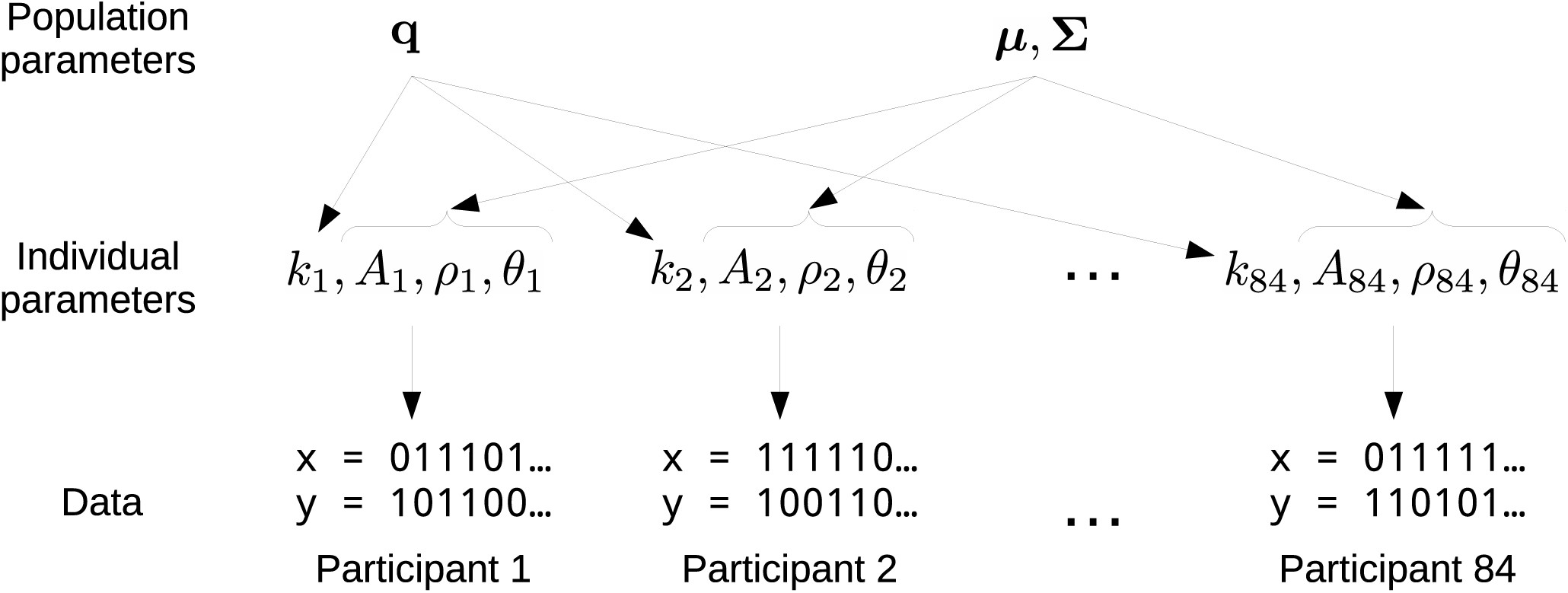
Hierarchical MPL model parameters. For each participant *i*, four parameters are fitted to the data: (*k_i_, A_i_, ρ_i_, θ_i_*). The population parameter *q* tracks the frequency of *k* values within the population, and the population parameters *µ* and Σ track the mean and covariance of (logit(*A*), logit(*ρ*), log(*θ*)) values within the population. The hierarchical PVL model differs from the MPL model by not having the *k* and *ρ* individual parameters and the *q* population parameter.

For each participant *i*, the WSLS model has four parameters 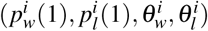. Vectors of transformed parameters 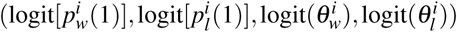 were given a multivariate Student’s *t* distribution with mean *µ*, covariance matrix Σ, and four degrees of freedom (*ν* = 4). The model’s hyperparameters were given weakly informative prior distributions. Each component of *µ* was given a normal prior distribution with mean 0 and variance 25, and Σ was decomposed into a diagonal matrix *τ*, whose diagonal components were given a half-normal prior distribution with mean 0 and variance 1, and a correlation matrix Ω, which was given an LKJ prior with shape *ν* = 1.

In summary, the hierarchical WSLS model is:

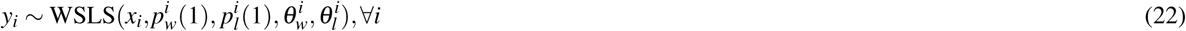

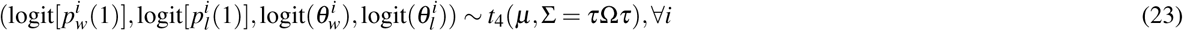

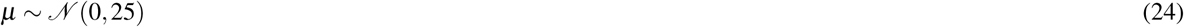

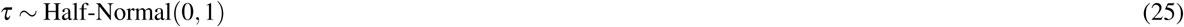

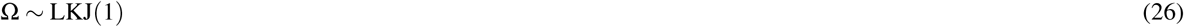

### Model fitting

Both models were coded in the Stan modelling language^59, 60^ and fitted to the data using the PyStan interface^61^ to obtain samples from the posterior distribution of model parameters. Convergence was indicated by 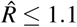 for all parameters, and at least 10 independent samples per chain were obtained^62^. All simulations were run at least twice to check for replicability.

### Model comparison

The PVL model includes parameters for learning decay and exploration to explain the participants’ behaviour in the probability learning task. The MPL model additionally includes parameters for pattern search and recency. We determined if pattern search and recency were relevant additions that increased the MPL model’s predictive accuracy (its ability to predict future data accurately) by comparing the PVL and MPL models using cross-validation. Additionally, we compared the PVL and MPL models to the WSLS model by the same method. (Since the CAB-*k* model^13^ is not a statistical model, it cannot be compared to the other models using cross-validation and for this reason has not been included in our model comparison.)

Statistical models that are fitted to data and summarised by a single point, their maximum likelihood estimates, can be compared for predictive accuracy using the Akaike information criterion (AIC). In this study, however, the three models were fitted to the data using Bayesian computation and many points of their posterior distributions were obtained, which informed us not only of the best fitting parameters but also of the uncertainty in parameter estimation. It would thus be desirable to use all the available points in model comparison rather than a single one. Moreover, the AIC’s correction for the number of parameters tends to overestimate overfitting in hierarchical models^62^. Another popular criterion for model comparison is the Bayesian information criterion (BIC), but it has the different aim of estimating the data’s marginal probability density rather than the model’s predictive accuracy^62^.

We first tried to compare the models using WAIC (Watanabe-Akaike information criterion) and the PSIS-LOO approximation to leave-one-out cross-validation, which estimate predictive accuracy and use the entire posterior distribution^66^, but the loo R package with which we performed the comparison issued a diagnostic warning that the results were likely to have large errors. We then used twelve-fold cross-validation, which is a more computationally intensive, but often more reliable, method to estimate a model’s predictive accuracy^66^. Our sample of 84 participants was partitioned into twelve subsets of seven participants and each model was fitted to each subsample of 77 participants obtained by excluding one of the seven-participant subset from the overall sample. One chain of 2,000 samples (warmup 1,000) was obtained for each PVL model fit, one chain of 5,000 samples (warmup 2,500) was obtained for each WSLS model fit, and one chain of 20,000 samples (warmup 10,000) was obtained for each MPL model fit (as the MPL model converges much more slowly than the other models). The results of each fit were then used to predict the results from the excluded participants as follows.

For each participant, 1,000 samples were randomly selected from the model’s posterior distribution and for each sample a random parameter set *ϕ* (e.g., *ϕ* = (*A, θ*) for the PVL model) was generated from the hyperparameter distribution specified by the sample. The probability of the *i*th participant’s results Pr(*y_i_|x_i_*) was estimated as

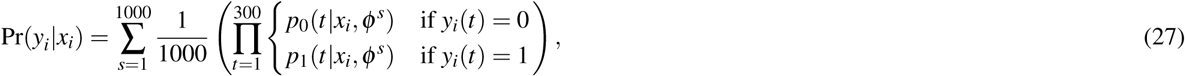

where 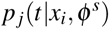 is the probability that the participant would choose option *j* in trial *t*, as predicted by the model with parameters *φ^s^*. The model’s estimated out-of-sample predictive accuracy CV was given by

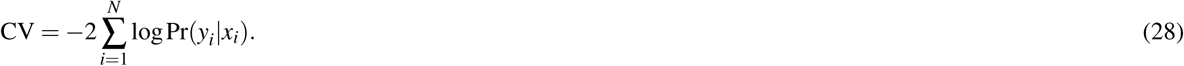

A lower CV indicates a higher predictive accuracy. This procedure was repeated twice to check for replicability.

### Posterior predictive distributions

We also simulated the MPL model to check its ability to replicate relevant aspects of the experimental data and predict the results of hypothetical experiments. To this end, two chains of 70,000 samples (warmup 10,000) were obtained from the model’s posterior distribution, given the observed behavioural data. A sample was then repeatedly selected from the posterior distribution of the hyperparameters (the population parameters *µ*, Σ, and *q*), random (*k,A,ρ,θ*) vectors were generated from the distribution specified by the sample, and the MPL model was simulated to obtain replicated prediction sequences *y* using the generated parameters on either random outcome sequences *x*, Pr(*x*(*t*) = 1) = 0.7, or the same *x* sequences our participants were asked to predict. By generating many replicated data, we could estimate the posterior predictive distribution of relevant random variables^62^. For instance, would participants maximise if they stopped searching for patterns? To answer this question, we simulated the model with *k* = 0 and (*A,ρ,θ*) randomly drawn from the posterior distribution, and calculated the mean response. If the mean response was close to 1, the model predicted maximisation.

### Data availability

All experimental data and computer code generated during and/or analysed during the current study are available at https://github.com/carolfs/m

## Results

### Behavioural results

For each trial *t*, we calculated the participants’ mean response, equal to the frequency of choice of the majority option. Results are shown in Fig. 3. Initially, the mean response was around 0.5, but it promptly increased, indicating that participants learned to choose the majority option more often than the minority option. The line *y* = 0.7 in Fig. 3 is the expected response for probability matching. In the last 100 trials of the task, the mean response curve is generally above probability matching: participants chose the majority outcome with an average frequency of 0.77 (*SD* = 0.10). The mean response in the last 100 trials was distributed among the 84 participants as shown in Fig. 4 (observed distribution).

**Figure 3.**
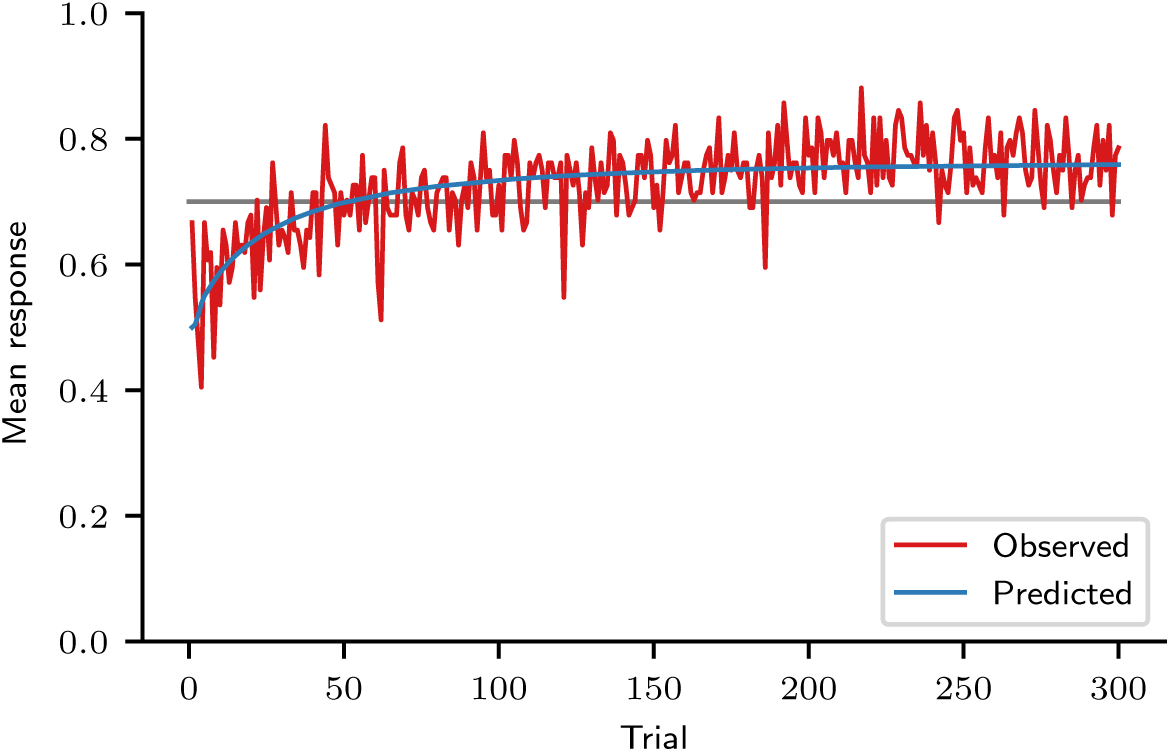
Mean response curve. Observed mean response curve of participants and predicted mean response curve, obtained by fitting the MPL model to the experimental data. The line *y* = 0.7 corresponds to the mean response of an agent that matches probabilities. (Participants: *N* = 84; MPL simulations: *N* = 10^6^.)

**Figure 4.**
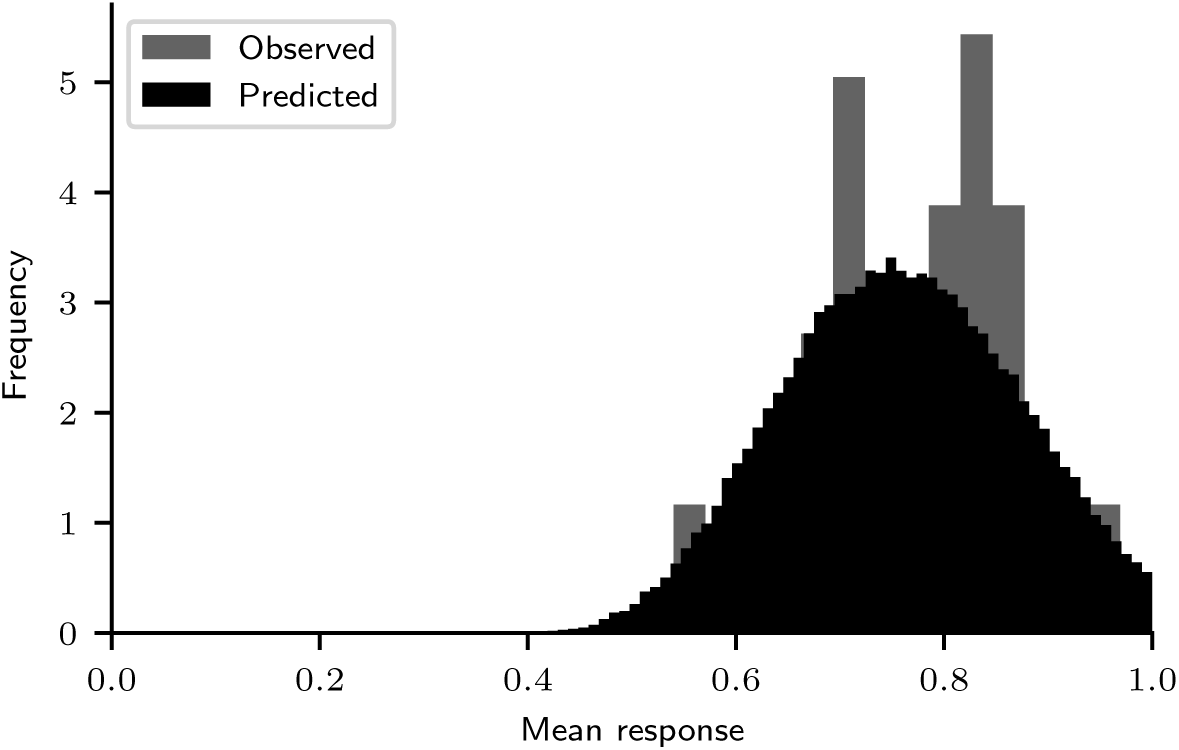
Predictive and observed mean response distributions in trials 200–300. (Participants: *N* = 84; MPL simulations: *N* = 10^5^.)

The cross-correlation of all participants was calculated for the last 100 trials, because in this trial range their mean response was relatively constant, as evidenced by Fig. 3. The average cross-correlation was 0.30 (*SD* = 0.19), implying that, on average, 65% of the participants’ predictions were equal to the previous outcome and consistent with the “win-stay, lose-shift” strategy. This cross-correlation value, however, can also be produced by pattern search strategies, as shown in Section “MPL model check: cross-correlation” below.

The nonmonotonic recency effect analysis results are shown in Fig. 5. They suggest that the nonmonotonic recency pattern, a component of the wavy recency effect that can be generated by pattern search, is present in trials 1–100, but not in trials 201–300. In the former trials, the mean response increased for three trials after a 0, decreased in the fourth trial, and increased again in subsequent trials. In the latter trials, after a 0 outcome, the mean response always increased. It stayed below the mean for the two subsequent trials after 0, indicating that participants predicted 0 at an above-average frequency in those two trials. From the third trial on, the mean response increased above the mean, indicating that participants predicted 0 at a below-average frequency. According to Plonsky et al.^13^, this result indicates that *k* = 3 in the first 100 trials, because the mean response curve is predicted to decrease in trial *k* + 1 after a 0 outcome. Indeed, a nonmonotonic recency pattern similar to the one observed in the first 100 trials can be obtained by simulating the MPL model with *k* = 3, *A* = 1, *ρ* = 1, and *θ →* ∞, which makes it equivalent to the CAB-*k* model with *k* = 3^13^ (Fig. 6). Alternatively, the observed pattern can be explained by expectation matching: since the probability that *x* = 0 is 0.3, four trials after the last 0 outcome is when one would expect the next 0 outcome to occur if 0 outcomes occurred regularly every four trials, with 1/4 = 0.25 probability. This would also explain why this pattern is only present in the first 100 outcomes: as responses are reinforced, participants make more habitual choices driven by reinforcement learning and fewer choices driven by cognitive biases such as expectation matching.

**Figure 5.**
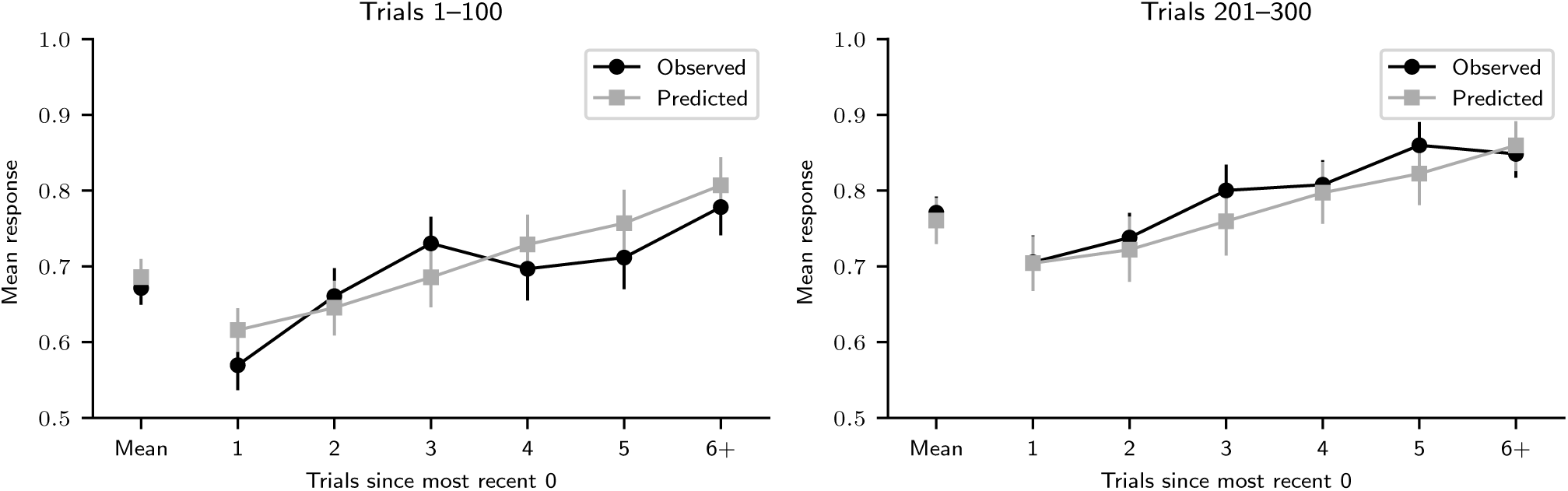
Nonmonotonic recency effect results. Nonmonotonic recency effect analysis results in trials 1–100 and 201–300 for observed data and predicted data, obtained by fitting the MPL model to the observed data. (Participants: *N* = 84; MPL simulations: *N* = 10^5^. The mean number of observations per participant or simulated agent for points 1 to 5 was 16.3 and for point 6+ was 16.5. The error bars are the 95% HDI.)

**Figure 6.**
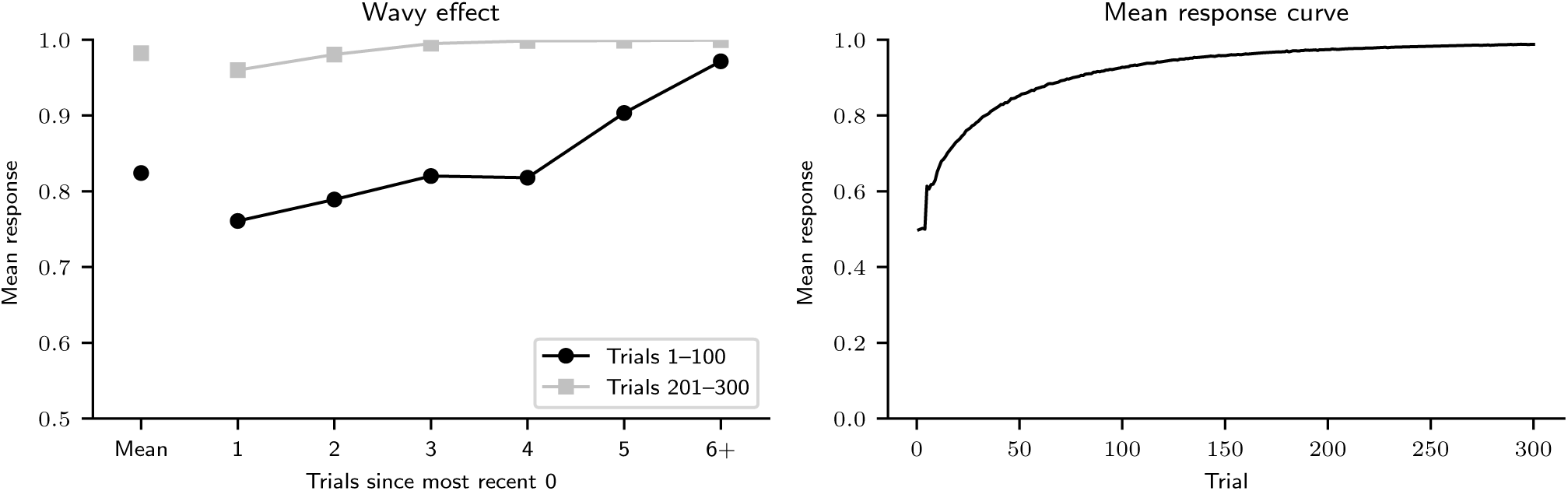
Nonmonotonic recency effect results for *k* = 3. Nonmonotonic recency effect analysis results in trials 1–100 and 201–300 (left) and mean response curve (right) for MPL agents with parameters *k* = 3, *A* = 1, *ρ* = 1, and *θ →* ∞ (*N* = 10^5^).

### Pattern learning by MPL agents

In this study we analysed behavioural data with the MPL model, a reinforcement model that searches for patterns. In the task we employed, however, participants were asked to predict outcomes that did not follow a pattern. To demonstrate how the MPL model learns patterns, thus, we must simulate MPL agents performing a different task. In this section we show that the MPL model with appropriate parameters can learn any pattern generated by a Markov chain of any order *L* ≥ 0. This includes all deterministic patterns, such as the repeating pattern 001010001100, of length 12, employed in a previous study with human participants^9^.

When the sequence to be predicted is generated by a fixed binary Markov chain of order *L*, the optimal strategy is to always choose the most likely outcome after each history *η* of length *L*. If an MPL agent is created with parameters *k ≥ L*, *A* = 1 (no forgetting), *ρ* = 1 (no recency), and *θ* → ∞ (no exploration), it will eventually learn the optimal strategy by the following argument. In this scenario, each expected utility will simply be a count of how many times that option was observed after the respective history, and the most frequent option will be observed more often than the least frequent one in the long run, which will eventually make its expected utility the highest of the two. The option with the highest expected utility will then be chosen every time, because this agent does not explore. If *k ≥ L*, the highest possible values for *A* (*A* = 1) and *θ* (*θ* → ∞) maximise the agent’s expected accuracy. A high *A* value means that past observations are not forgotten, which is optimal, because the Markov transition matrix that generates the sequence of outcomes is fixed and past observations represent relevant information. In this task, exploration, i.e. making random choices due to *θ <* ∞, does not uncover new information, because the agent always learns the outcomes of both options, regardless of what it actually chose. Thus, a high *θ* value is optimal, as it means that the “greedy” choice (of the option with the highest expected utility) will always be made.

Table 1 demonstrates how two MPL agents learn a deterministic alternating pattern in an eight-trial task. First, note that an alternating sequence, 01010101…, is formed by repeating the subsequence 01 of length 2, but can be generated by a Markov chain of order 1, where 0 transitions to 1 with 1 probability and 1 transitions to 0 with 1 probability. The MPL agent therefore only needs *k* = 1 to learn it, and only needs to consider two histories of past outcomes: *η* = 0 and *η* = 1. Similarly, the repeating pattern 001010001100 of length 12^9^ can be generated by a Markov chain of order 5, and an MPL agent only needs *k* = 5 to learn it. (These grammar rules generate the pattern 001010001100: 00101 → 0, 01010 → 0, 10100 → 0, 01000 → 1, → 10001 → 1, 00011 → 0, 00110 → 0, 01100 → 0, 11000 → 0, 10000 → 1, 00001 → 0, 00010 → 1. They prove that the pattern can be generated by a Markov chain of order 5.)

The left half of Table 1 demonstrates how an agent with optimal parameters for this task (*k* = 1, *A* = 1, *ρ* = 1, *θ →* ∞) learns the pattern. Initially, in trial *t* = 1, the expected utilities of predicting 0 or 1 are 0 for both histories *η* = 0 and *η* = 1. Similarly, in trial *t* = 2, a history of length 1 has not yet been observed, and the agent just predicts 0 or 1 with 0.5 probability (*p*_1_ = 0.5). The outcome in trial *t* = 1 is *x* = 0, the first element of the alternating pattern. In trial *t* = 2, the agent has observed the history *η* = 0, but it has not learned anything about it yet and thus predicts 0 or 1 with 0.5 probability. It then observes that the outcome alternates to *x* = 1 and updates the expected utility of making a prediction after 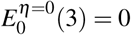 and 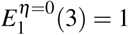. Thus, alternating to 1 after 0 acquires a higher expected utility than repeating 0 after 0. Since *A* = 1 and *ρ* = 1, this knowledge will not decay, and since *θ* → ∞, the agent will always exploit and predict 1 after 0. It has thus already learned half of the pattern. In trial *t* = 3, the agent has observed history *η* = 1, but it has not learned anything about it yet and thus predicts 0 or 1 with 0.5 probability. It then observes that the outcome is *x* = 0 and updates the expected utility of making a prediction after 1: 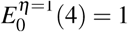 and 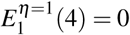. Since *A* = 1 and *ρ* = 1, this knowledge will not decay, and since *θ →* ∞, the agent will always exploit and predict 0 after 1. It has thus learned the entire pattern, and from trial *t* = 4 on it will always predict the next outcome correctly. In this example, the 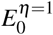 and 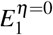 values count how many times the agent has observed 0 after 1 and 1 after 0 respectively.

The right half of Table 1 demonstrates how an agent with suboptimal parameters for this task (*k* = 1, *A* = 0.9, *ρ* = 0.9, *θ* = 0.3) also learns the pattern, but does not always make the correct prediction. Note that the 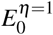 and 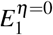 values decrease if the respective history has not been observed, as *A* = 0.9, and that even if the history is observed, the expected utility value increases by less than one, because *Aρ* = 0.81. Despite the learning decay the agent experiences, though, by *t* = 4, it has also learned the alternating pattern. If *θ* → ∞, it would always exploit and make correct predictions, but since *θ* = 0.3, it will frequently, but not always, make the correct prediction, as shown by the *p*_1_ column.

Fig. 7 shows the results of simulations wherein MPL agents with *A* = 1, *ρ* = 1, *θ* → ∞, and *k* = 0, 1, 2, 3 attempt to learn patterns of increasing complexity in a 300 trial task. An alternating pattern (left graph of Fig. 7) cannot be learned by an agent with *k* = 0. Agents with *k* ≥ 1 can learn the pattern, as demonstrated by their perfect accuracy in the last 100 trials of the task, even though learning this pattern only requires *k* = 1. In general, when *k < L*, the MPL model does not always learn the optimal strategy. The pattern 0011, of length 4, can be learned by agents with *k* ≥ 2 (middle graph of Fig. 7), and the pattern 110010, of length 6, by agents with *k* ≥ 3 (right graph of Fig. 7). These results again demonstrate that an agent with working memory usage *k* may be able to learn patterns of length greater than *k*.

**Figure 7.**
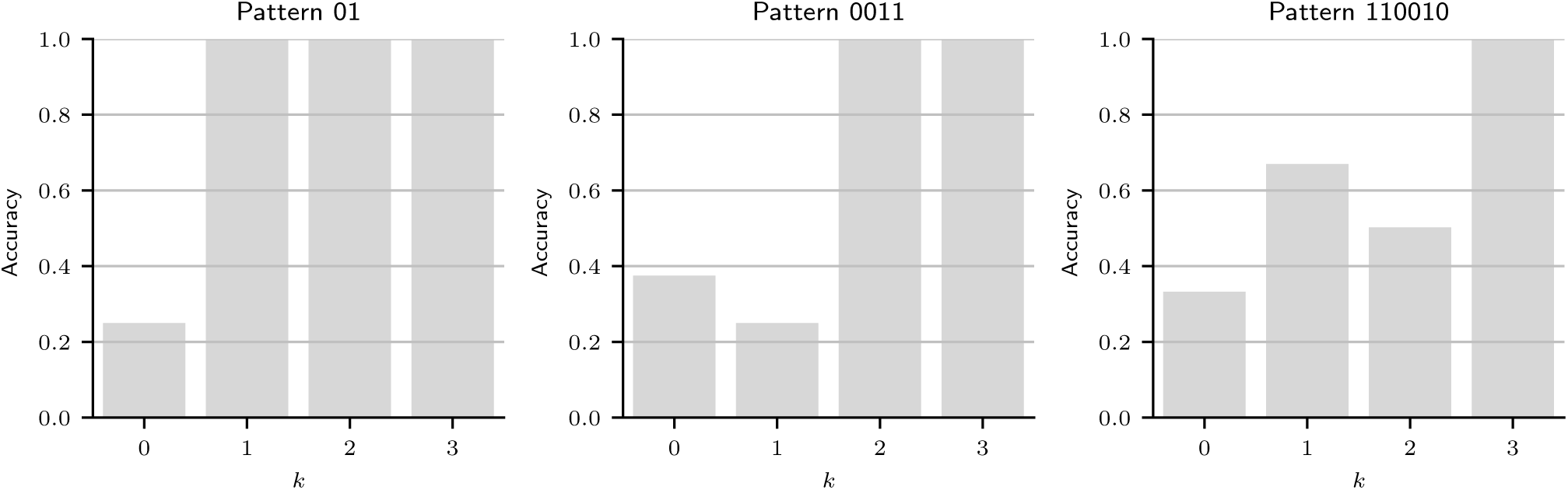
Accuracy of MPL agents in a pattern search task. Accuracy of MPL agents with varying working memory usage (*k*), *A* = 1, *ρ* = 1, and *θ →* ∞ in the last 100 of 300 trials for three different tasks, whose outcomes were generated by repeating the binary pattern strings 01, 0011, or 110010.

#### MPL parameter recovery

For each of 10,000 sets of random MPL parameters, an MPL agent was simulated. The MPL agent performed the same task as our participants, and the results were analysed using a Bayesian model. Figure 8 shows the Kullback–Leibler divergence between the model’s prior and posterior distributions of model parameters. This measures how much information about the parameters could be recovered from each agent’s results. When the Kullback–Leibler divergence is zero, no information is gained by running the analysis. The Kullback–Leibler divergence is given as a function of the agent’s *θ*, the exploration parameter, because when *θ* = 0, choices are completely random and thus the data contain no information about *A*, *ρ*, and *k*. Moreover, if *k* = 0, it is not possible to recover information about the *A* and *ρ* parameters separately, only about their product; thus, if the agent had *k* = 0, the Kullback–Leibler divergence was calculated for *Aρ*.

**Figure 8.**
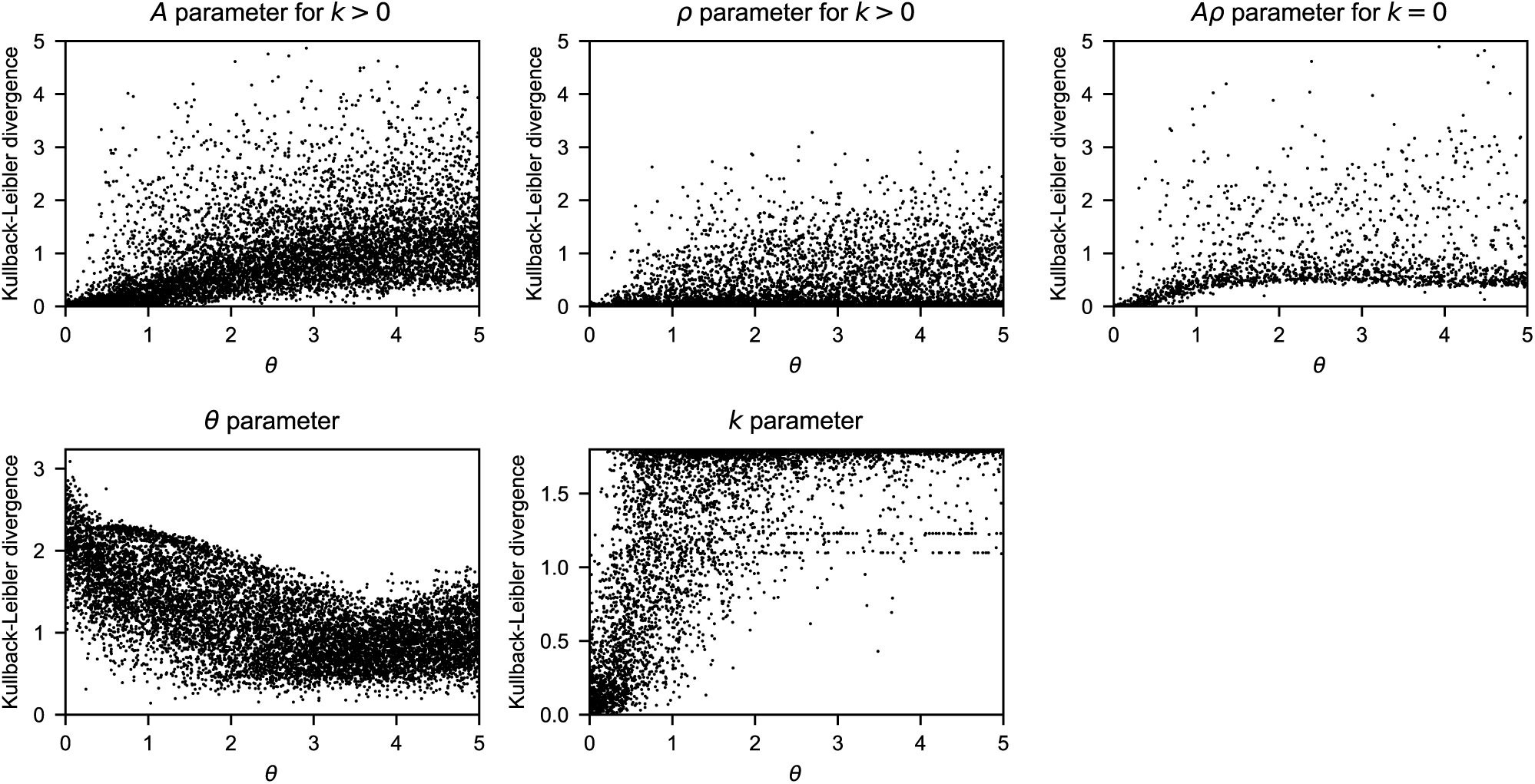
MPL parameter recovery from simulated data. The Kullback-Leibler divergence between the prior and posterior distributions is given for the parameters *A*, *ρ*, *Aρ*, *k*, and *θ* as a function of *θ*. The divergence was calculated separately for *A* and *ρ* if *k >* 0 and for the product *Aρ* if *k* = 0.

In the section “Posterior distribution of MPL model parameters” below, it was found that our human participants had a median *θ* = 0.23 and about 50% of predicted *θ* values were in the [0.15,0.37] interval. In this interval, the mean Kullback–Leibler divergences for *A*, *ρ*, and *Aρ* were 0.09, 0.03, and 0.08 respectively. For reference, the divergence between the uniform distribution and the beta distribution with *α* = 2, *β* = 2 is ≈ 0.125, greater than the observed values, and the latter distribution is very broad, with a 95% HDI of [0.094, 0.906]. Moreover, in this *θ* interval, the mean Kullback–Leibler divergence for *k* was 0.43, and for reference a single *k* can only be identified with 0.75 probability or greater if the divergence is at least 0.827, and it can only be identified with certainty if the divergence is 1.79. These results demonstrate that if *θ* is small, only very little can be learned from a single agent’s sequences about the *A*, *ρ*, and *k* parameters, and we would not be able to learn much from analysing each of our participants’ results in isolation. Hence, in this study, we applied a hierarchical model, which is fitted to the entire data set simultaneouly. The hierarchical model, in stark contrast with the individual model just discussed, allowed us to obtain much more precise results about the population distribution of MPL parameters (see section “Posterior distribution of MPL model parameters” below).

### Model comparison

The PVL, MPL, and WSLS models were compared by cross-validation. The PVL model obtained a cross-validation score of 2.731 × 10^4^, the MPL model obtained a cross-validation score of 2.656 × 10^4^, and the WSLS obtained a cross-validation score of 2.980 × 10^4^. The lower score for the MPL model suggests that the MPL model has a higher predictive accuracy than the PVL and WSLS models and thus that reinforcement learning and pattern search increased the MPL model’s ability to predict the participants’ behaviour. It also supports our use of the MPL model to predict the results of hypothetical experiments.

### Posterior distribution of MPL model parameters

Figs. 9 and 10 show the marginal posterior distributions of the parameters *k*, *A*, *B*, and *θ*. The most frequent *k* values were 0, 1, and 2, whose posterior probabilities were 0.15 (95% HDI [0.06,0.24]), 0.39 (95% HDI [0.25,0.53]), and 0.45 (95% HDI [0.32,0.59]) respectively. The posterior probability that *k* = 1 or *k* = 2 was 0.84 (95% HDI [0.75,0.93]), the posterior probability that *k* ≥ 1 (i.e., the participant searched for patterns) was 0.85 (95% HDI [0.76, 0.94]), and the posterior probability that *k* ≥ 3 was 0.01 (50% HDI [0.00,0.00], 95% HDI [0.00,0.06]). The posterior medians of *A, ρ*, and *θ*, given by the transformed *µ* parameter, were 0.99 (95% HDI [0.98, 0.99]), 0.96 (95% HDI [0.95,0.98]), and 0.23 (95% HDI [0.19,0.28]) respectively. The posterior distribution of the correlation matrix indicates that the correlation between *A* and *ρ* is 0.14 (95% HDI [0.34,0.61]), the correlation between *A* and *θ* is −0.56 (95% HDI [−0.87,0.20]), and the correlation between *ρ* and *θ* is −0.76 (95% HDI [−0.91,0.57]).

**Figure 9.**
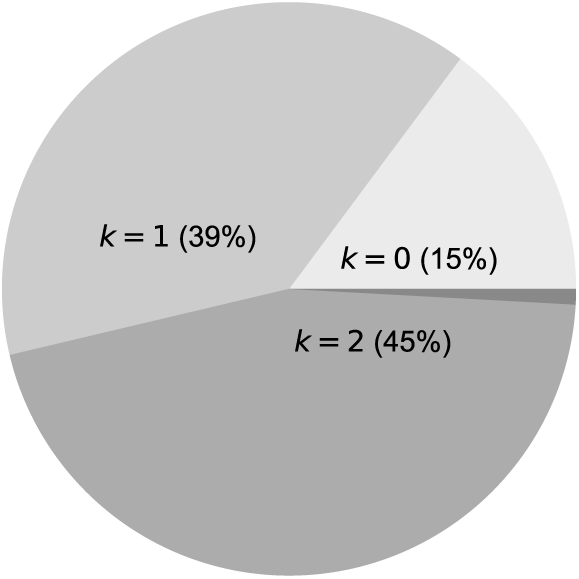
Marginal posterior distribution of *k*. It is given by the mean of the *q* parameter (see Fig. 2).

**Figure 10.**
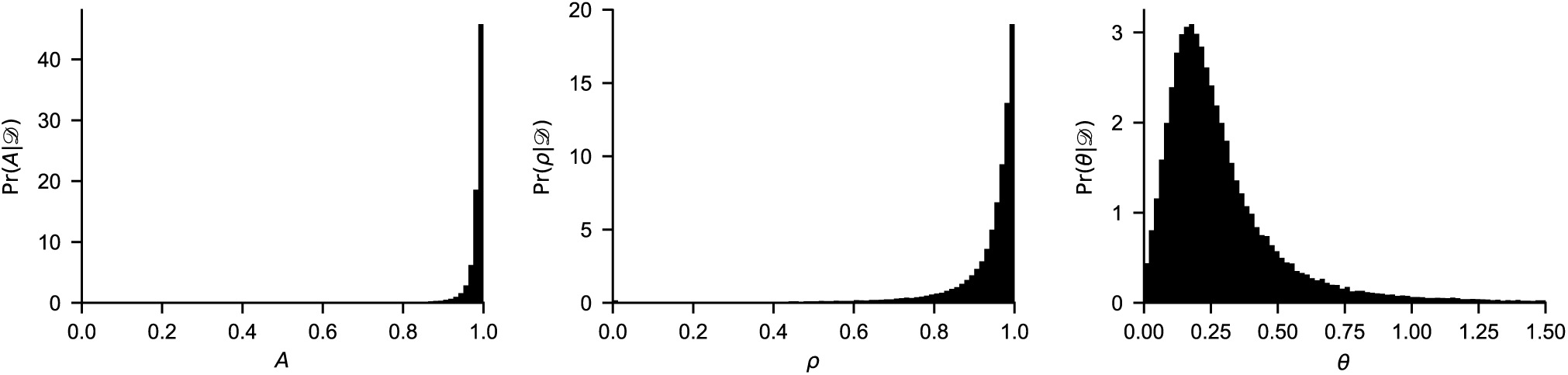
Marginal posterior distributions of *A*, *B*, and *θ*, given the observed data *D*. The graphs were obtained by generating random (*A, ρ, θ*) vectors from the posterior distribution of model hyperparameters.

Although the posterior medians of *A* and *ρ* were very close to 1, their upper limit, these values still imply significant recency and forgetting, because the effect of *A* and *ρ* is exponential. A value of 0.95, for instance, implies that participants would forget nearly all (~92%) of what they had learned before the last 50 trials, because 0.95^50^ = 0.08. Even a value of 0.99 still implies an information loss of 40% within 50 trials.

### MPL model check: mean response

Fig. 3 displays the predicted mean response curve. The predicted mean response in the last 100 trials is 0.76 (95% HDI [0.54,0.96]) for a new participant and 0.76 (95% HDI [0.74,0.78]) for a new sample of 84 participants and the same *x* sequences our participants predicted. The latter prediction is consistent with the observed value: 11% of samples are predicted to have a mean response as high or higher than observed (0.77). The predicted standard deviation of the mean response in the last 100 trials for 84 participants is 0.11 (95% HDI [0.09, 0.13]), and 96% of samples are predicted to have a standard deviation as high or higher than observed (0.10). The predicted and observed mean response distributions are shown in Fig. 4.

### MPL model check: cross-correlation

As previously discussed, a “win-stay, lose-shift” behaviour can be generated by the MPL model with *k* = 0 and *Aρ* = 0. However, the posterior distribution of parameters we obtained suggests the opposite of “win-stay, lose-shift:” *k* is greater than 0 with 0.85 probability and the medians of *A* and *ρ* are close to 1. Even though the MPL model had a better cross-validation score than the WSLS model, since previous studies that suggest many participants use a “win-stay, lose-shift” strategy^9, 39^, this raises the possibility that our analysis is not consistent with the experimental data. To check for this possibility, we calculated the predicted cross-correlation *c*(*x,y*) between *y* and *x* in the last 100 trials of the task.

The predicted cross-correlation for a new sample of 84 participants performing the task with the same *x* sequences was 0.28 (95% HDI [0.25, 0.32]), and 10% of participant samples are predicted to have an average cross-correlation as high or higher than observed (0.30). The observed cross-correlation is thus consistent with what MPL model predicts, suggesting that it does not reflect a “win-stay, lose-shift” strategy; rather, this result indicates that most participants adopted a pattern-search strategy, which also produced many responses that were incidentally equal to the previous outcome.

### MPL model check: nonmonotonic recency effect

Fig. 5 displays the predicted mean response as a function of trials since the most recent *x* = 0 outcome, generated by simulating MPL agents with parameters randomly drawn from the posterior distribution, performing the probability learning task with the same *x* sequences as our participants. The predicted mean response trend, both for the first and the last 100 trials, is increasing rather than wavy. The model thus predicts the observed trend accurately in the last 100 trials, but not in the first 100 trials. This is consistent with the explanation that the nonmonotonic pattern observed in the first 100 trials is due to expectation matching rather than pattern search. If expectation matching strongly influenced the participants’ choices in the first trial range but not in the last one, the MPL model would only be able to predict the results accurately in the latter, since it does not implement expectation matching.

### Predicted effect of outcome probabilities

Both the observed and predicted mean responses in the last 100 trials, 0.77 and 0.76 respectively, approximately matched the majority outcome’s probability, 0.7. Would the MPL model also predict probability matching for a new sample of participants if the outcome probabilities were different? Fig. 11 shows the mean response curve for different values of the majority outcome’s probability *p*, as predicted by the MPL model with parameters fitted to our participants. The predicted mean response increased with *p*. If *p* = 0.5,0.6*,…,* 1.0, the predicted mean responses at *t* = 1000 were 0.50, 0.65, 0.76, 0.85, 0.90, and 0.96 respectively. Thus, the MPL model with fitted parameters predict approximate probability matching.

**Figure 11.**
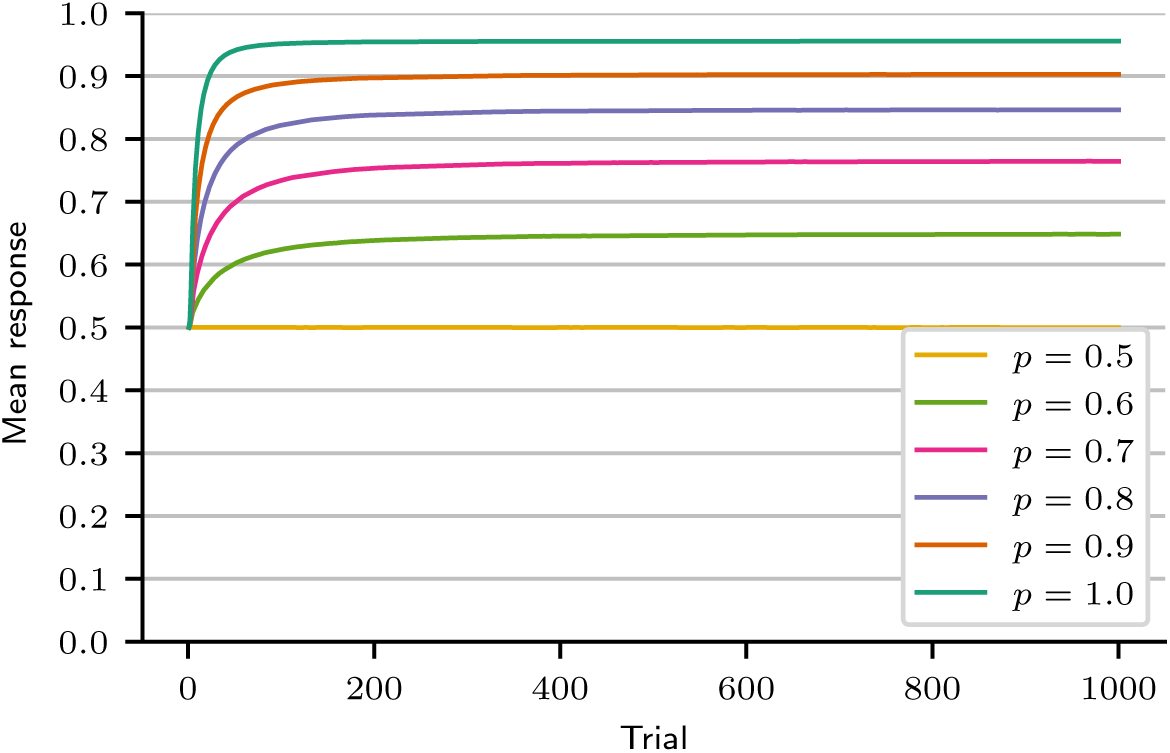
The predicted mean response increases with the probability of the majority option (*p*). Results were obtained by simulation using the posterior distribution of MPL model parameters. (*N* = 10^6^ by *p* value.)

### Predicted effect of pattern search, exploration, and recency on learning speed and mean response

As demonstrated in Section “Pattern learning by MPL agents”, an MPL agent performs optimally in a task without patterns if *k* = 0 (no pattern search), *A* = 1 (no forgetting), *ρ* = 1 (no recency), and *θ* → ∞ (no exploration). Other parameter values, however, do not necessarily lead to a suboptimal performance. In particular, an agent that searches for patterns (*k >* 0) may also maximise. This is shown in the top left graph of Fig. 12. If *A* = 1, *ρ* = 1, and *θ* → ∞, the mean response eventually reaches 1 (maximisation) even if *k >* 0. In fact, as shown in the top right graph of Fig. 12, agents will learn to maximise even if *θ* = 0.3, which is approximately the median value estimated for our participants. If *A <* 1, however, agents that search for patterns never learn to maximise, as the bottom left graph of Fig. 12 demonstrates. And if *ρ <* 1, no agent learns to maximise, as the bottom right graph of Fig. 12 demonstrates. Thus, pattern search only decreases long-term performance compared to no pattern search when combined with forgetting. As *k* increases, however, pattern-searching agents take longer to maximise, especially if *θ* is low. The MPL model thus suggests that pattern search impairs performance by slowing down learning in the short term (top left graph of Fig. 12) and, when combined with forgetting, in the long term (bottom left graph of Fig. 12). The former has already been proposed by Plonsky et al.^13^ using other models of pattern search.

**Figure 12.**
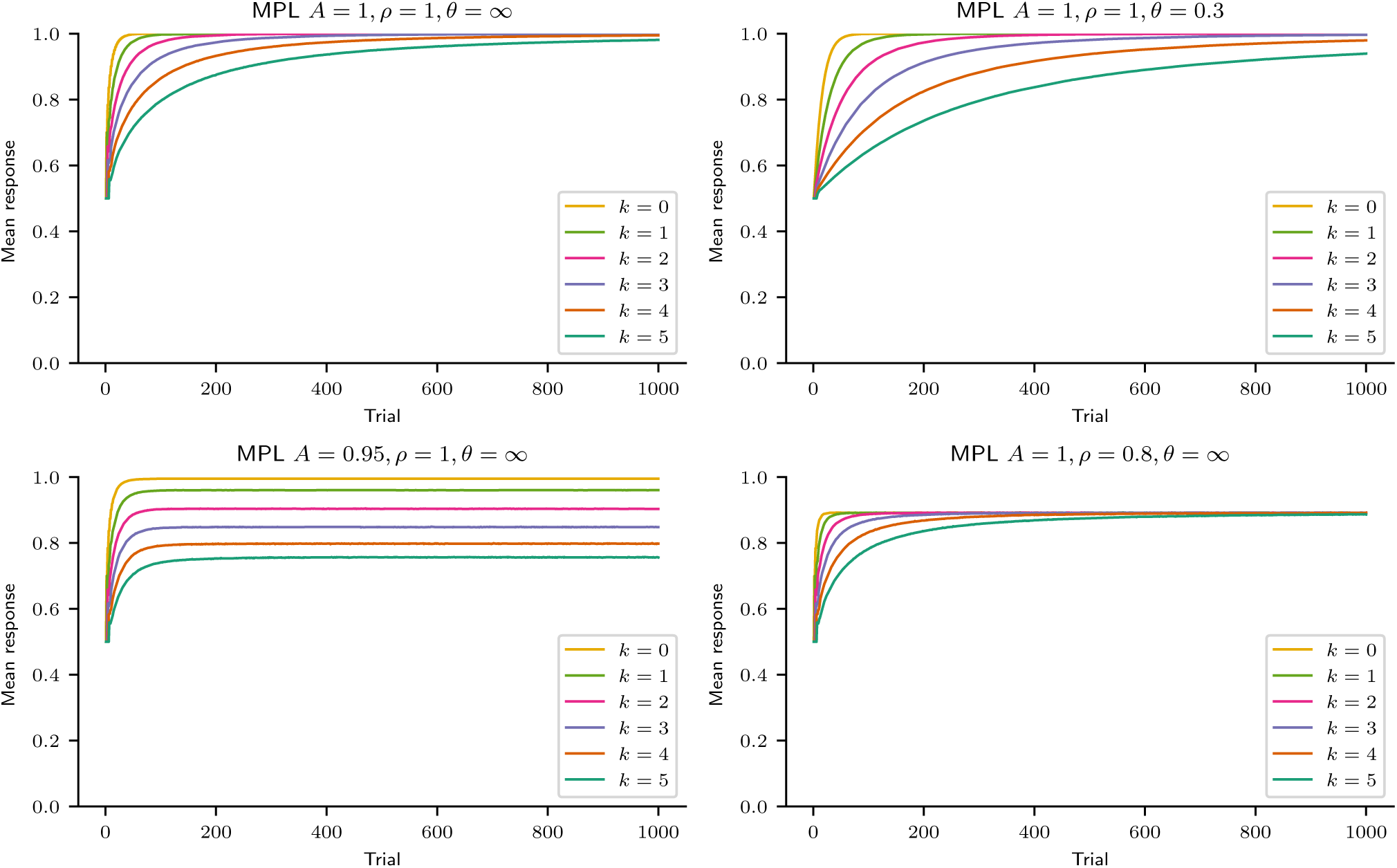
Mean response curve of MPL model agents performing a probability learning task. Simulations of the MPL model indicate that pattern search (*k >* 0) does not necessarily decrease the asymptotic mean response in a 1000-trial probability learning task, but agents who search for patterns are slower to learn the majority option (top). Pattern search combined with forgetting (*k >* 0, *A <* 1), as well as recency (*ρ <* 1), decreases the asymptotic mean response (bottom). (*N* = 10^6^ by parameter set.)

The bottom row of Fig. 12 also demonstrates that the parameters *A* and *ρ* have distinct effects on performance if *k >* 0. If *A <* 1 (forgetting occurs, left graph), agents perform worse and worse as the complexity of pattern search increases, but if *ρ <* 1 (recency occurs, right graph), agents never perform optimally and their asymptotic performance does not depend on pattern search complexity.

How much did pattern search actually affect our participants’ performance, though? Fig. 13 shows the predicted mean response curve for participants with *k* from 0 to 3, using the obtained posterior distribution of *A*, *ρ*, and *θ*. Participants with low *k* are expected to perform better than participants with high *k*, especially in the beginning, although, since *ρ <* 1, even participants with *k* = 0 (no pattern search) should not maximise. In the last 100 of 300 trials, a participant with *k* = 0, 1, 2, 3 is predicted to have a mean response of 0.82 (95% HDI [0.60, 1.00]), 0.77 (95% HDI [0.56, 0.96]), 0.72 (95% HDI [0.52, 0.89]), and 0.67 (95% HDI [0.49, 0.82]) respectively. Note that the model predicts that mean response variability is high for each *k* and thus that *k* is a weak predictor of mean response.

**Figure 13.**
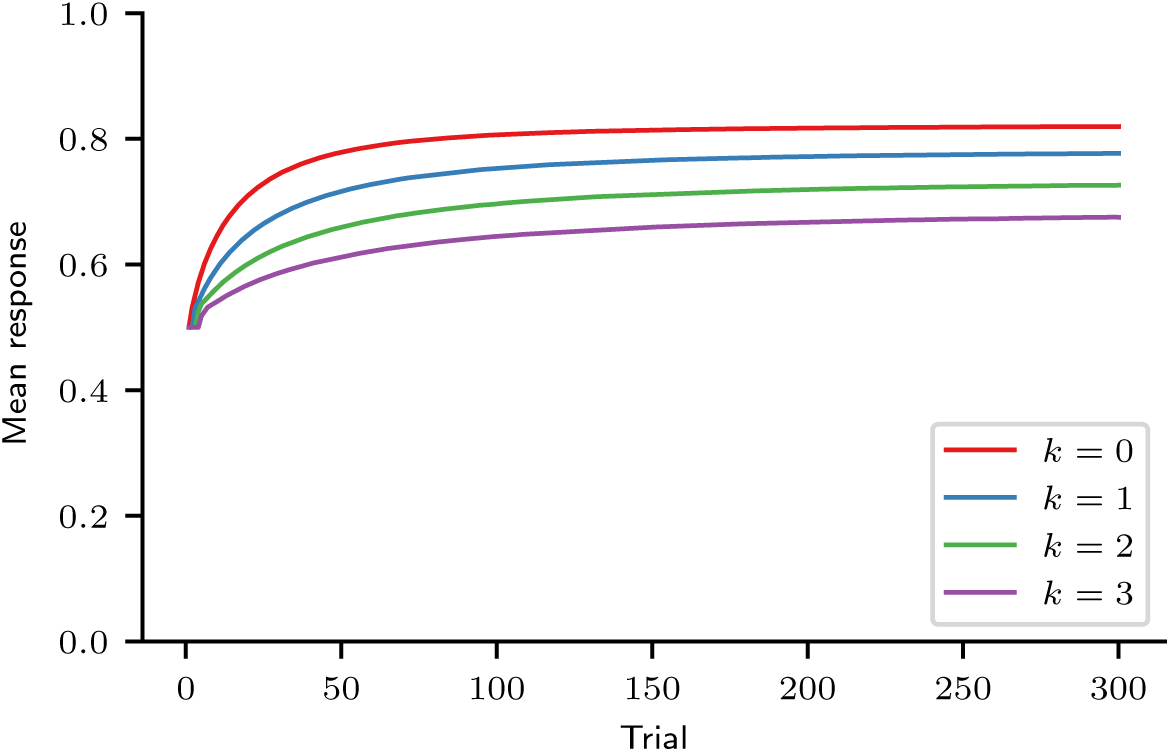
Predicted mean response curve for *k* = 0, 1, 2, 3. Results were obtained by simulation using the posterior distribution of MPL model parameters. (*N* = 10^6^ by *k* value.)

The difference between the *k* = 0 and *k* = 2 mean response curves is largest (0.11 on average) in the 100-trial range that spans trials 18-117. To check if this difference in mean response could be detected in our experimental results, a linear regression was performed in the logit scale between the participants’ mean *k* estimates and their observed mean responses in the trial ranges 18-117 and 201-300, using ordinary least squares. The results are shown in Fig. 14. In both trial ranges, the mean response decreased with the mean *k*, as indicated by the negative slopes, but in trials 201-300, as expected, this trend was smaller. Moreover, in both trial ranges the small *R*^2^ indicates that the mean *k* is a weak predictor of mean response.

**Figure 14.**
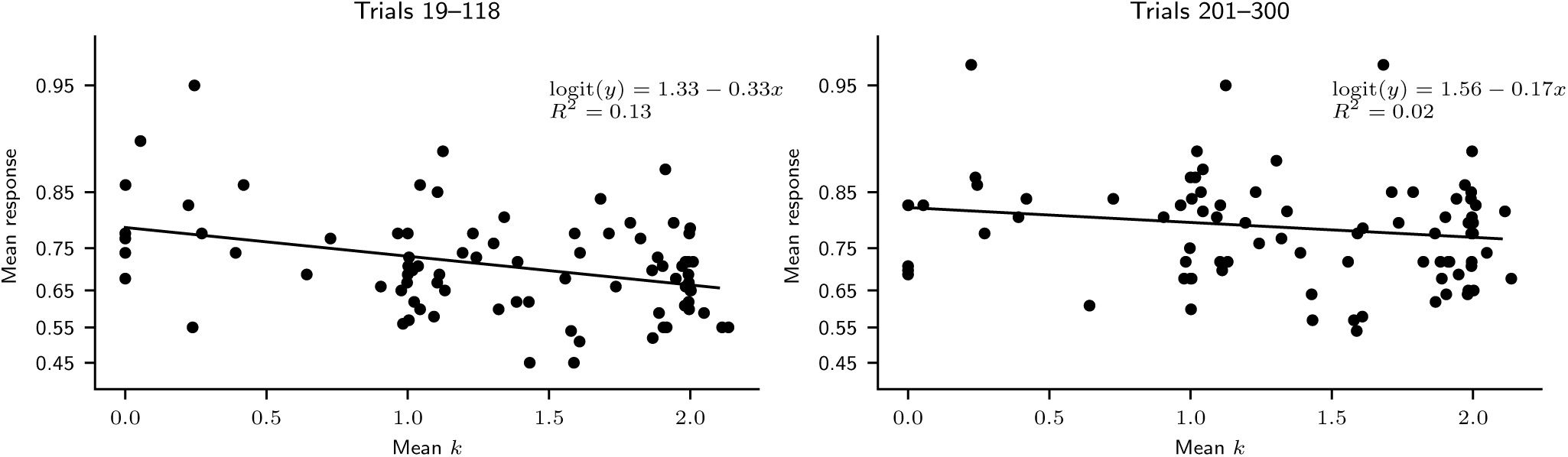
Mean response of participants in trials 18–117 (left) and 201–300 (right) as a function of their mean *k*. (*N* = 84.)

To predict the effect of pattern search (*k >* 0), exploration (*θ <* ∞), and recency (*ρ <* 1) on our participants’ performance, we simulated hypothetical experiments in which participants did not engage in one of those behaviours, using the MPL model with parameters fitted to our participants. We did not simulate an experiment in which participants did not forget what they had learned (*A* = 1) because we assume that forgetting was not affected by our participants’ beliefs and strategies. In the last 100 of 300 trials, the predicted mean response was 0.82 for a “no pattern search” experiment, 0.89 for a “no recency” experiment, and 0.94 for a “no exploration” experiment (Fig. 15). Thus, “no exploration” has the largest impact on mean response, followed by “no recency,” and lastly by “no pattern search.”

**Figure 15.**
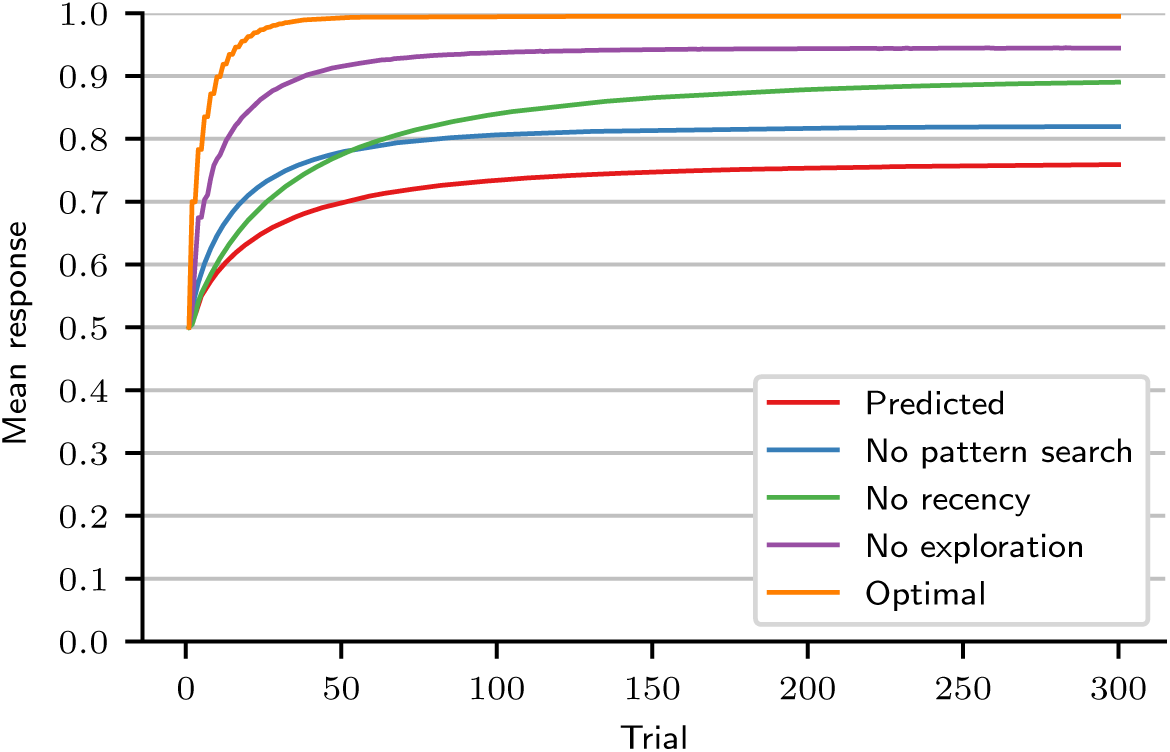
Mean response curve for a replication of this experiment (predicted) and for hypothetical experiments in which participants engaged in no pattern search, or no recency, or no exploration, or none of those behaviours (optimal). Results were obtained by simulation using the posterior distribution of MPL model parameters. (*N* = 10^6^ by curve.)

## Discussion

In this study, 84 young adults performed a probability learning task in which they were asked to repeatedly predict the next element of a binary sequence. The majority option had 0.7 probability of being rewarded, while the minority option had 0.3 probability of being rewarded. The optimal strategy—maximising—consisted of always choosing the majority option. Our participants chose that option in the last 100 of 300 trials with 0.77 frequency. This is consistent with numerous previous findings, which show that human participants generally do not maximise; instead, they approximately match probabilities^1–3^. Previous research also suggests that participants search for patterns in the outcome sequence^2, 5–11^. For this reason, we modelled our data with a reinforcement learning model that searches for patterns, the Markov pattern learning (MPL) model. In a model comparison using cross-validation, the MPL model had a higher predictive accuracy than the PVL and WSLS models, which do not search for patterns^37, 38^. This is additional evidence that participants indeed search for patterns. The fitted MPL model could also predict accurately all the features of the behavioural results in the last 100 trials that we examined: the participants’ mean response and mean response standard deviation, the cross-correlation between the sequences of outcomes and predictions, and the mean response as a function of the number of trials since the last minority outcome (the “nonmonotonic recency effect” analysis).

As discussed in the Introduction, the model does not estimate, and thus cannot explicitly match, the outcome probabilities; nevertheless its average behaviour, after being fitted to the data, approximately matched them, even in simulations in which the outcome probabilities were different from 0.7/0.3. Similarly, our human participants may not have been trying to match probabilities, even though they did. This justifies switching our focus from why participants matched probabilities to why they simply failed to perform optimally.

Our analysis indicates that 85% (95% HDI [76,94]) of participants searched for patterns and took into account one or two previous outcomes—*k* = 1 or *k* = 2—to predict the next one. This finding challenges the common claim that many participants use the “win-stay, lose-shift” strategy^9, 39^, since this strategy implies *k* = 0. In one study^9^, more than 30% of participants in one experiment and more than 50% of participants in another were classified as users of “win-stay, lose-shift.” Based on our analysis, we would claim instead that no more than 15% (95% HDI [6,24]) of participants (those with *k* = 0) could have used “win-stay, lose-shift.” We checked this claim by calculating the observed and predicted cross-correlations between the sequences of outcomes and predictions, since “win-stay, lose-shift” creates a high cross-correlation. The observed cross-correlation, which indicated that about two thirds of predictions were consistent with “win-stay, lose-shift,” was also consistent with what the MPL model predicted, providing evidence that our analysis is accurate and that pattern search can also produce the observed cross-correlation. This conclusion was further supported by the MPL model having a higher predictive accuracy than the WSLS model in a model comparison using cross-validation.

Our results, which suggest that *k* ≤ 2 for 99% of participants (95% HDI [94, 100]), also disagree with the results obtained by Plonsky et al.^13^, which suggest that participants performing a 100-trial reinforcement learning task employed much higher *k* values, such as *k* = 14. To check our results against those of Plonsky et al.^13^, we adapted to our study design the nonmonotonic recency effect analysis proposed by them. Our data set exhibited a nonmonotonic recency effect in the first 100 trials of the task, but not in the last 100 trials, where the mean response always increased after a loss. Simulated data using the MPL model with fitted parameters displayed an increasing trend instead of a nonmonotonic recency effect in both the first and the last 100 trials. If the interpretation of this pattern presented by Plonsky et al.^13^ is correct, i.e., it is caused by pattern search, then our data analysis indicates that *k* = 3 in the first 100 trials. Indeed, simulated MPL agents with *k* = 3 (equivalent to CAB-*k* agents with *k* = 3) did exhibit a nonmonotonic recency effect like the observed one. However, the same agents also maximised instead of matched probabilities. This is because while pattern search impairs performance, as demonstrated by Plonsky et al.^13^ and the present study, it is necessary to employ large *k* values such as *k* = 14 to impair performance to the level of probability matching. Thus, pattern search with *k* = 3 explains the nonmonotonic recency effect observed in the first 100 trials of the task, but it does not explain probability matching.

The same observations are, however, compatible with the alternative proposal that the wavy recency effect is caused by expectation matching. In this scenario, we would expect the lowest mean response to occur three to four trials after a loss, since the probability that *x* = 0 is 0.3. This was observed in the first 100 trials of the task, and explains why the MPL model with fitted parameters was not able to predict those results accurately—the model does not include expectation matching. As responses were reinforced along the task, participants would have learned to make more choices driven by reinforcement learning and fewer driven by expectation matching, which explains why the nonmonotonic recency effect was not found in the last 100 trials and why the MPL model with fitted parameters could predict those results accurately. We conclude that the nonmonotonic recency effect found in the first 100 trials does not contradict our analysis suggesting *k* ≤ 2. This estimate is also consistent with the estimated capacity of working memory (about four elements), while large *k* values such as *k* = 14, required to explain probability matching, are not^19^.

Our MPL simulations agree with the basic premise in Plonsky et al.^13^ that the search for complex patterns, employing large *k* values, leads to a suboptimal performance because of the “curse of dimensionality.” Since, however, participants seem to have searched only for simple patterns, the suboptimal performance observed in the last 100 trials could not have been caused by this effect. It might still have been caused, in principle, by the interaction between pattern with forgetting (Fig. 13). Because of forgetting, participants with *k* = 0, who do not search for patterns, are predicted to achieve a mean response in the last 100 trials 10% higher than participants with *k* = 2, and 6% above average. But this is only a small improvement. It indicates that even participants who did not search for patterns were on average still far from maximising. Indeed, in our experimental data, a lower mean *k* was associated with an only slightly higher mean response and mean *k* was a weak predictor of mean response. This suggests that pattern search is not the main behaviour that impairs performance, and that decreasing working memory usage for pattern search would not lead to maximising. Indeed, in a previous study, participants matched probabilities in a probability learning task whether or not their working memory was compromised in a dual-task condition^63^.

The main behaviours that decreased performance the most, according to our analysis, were exploration and recency. Exploration is adaptive in environments where agents can only learn an option’s utility by selecting it and observing the outcome. In our task, participants did not have to select an option to learn its utility; they could use fictive learning to do so. Nevertheless, our simulations suggest that participants did explore, and that if they had not explored, their mean response in the last 100 trials would increase by 19%. In comparison, if they had not searched for patterns, their mean response would increase by only 6%.

This conclusion does not necessarily contradict the pattern search hypothesis. Plonsky et al.^13^ and the current work define pattern search as the learning of relationships between an event and the events that preceded it. Exploration, on the other hand, is a tendency for choosing an option at random when both options have similar expected utilities. These definitions clearly distinguish pattern search from exploration in the present work. Nevertheless, when participants explore and choose an option that has not been previously reinforced, they may be trying to follow some pattern or rule they just thought up. According to our definitions, this behaviour is exploration, not pattern search, because it ignores learned relationships between events (and is thus rather inefficient at finding patterns), but it fits the more general view of pattern search put forward by Wolford et al.^6^. We do not know, however, the exact reasons behind exploratory choices. When participants select a random option, they may just be taking a guess rather than thinking about a pattern. In this work, we call “exploration” all behaviours not explained by pattern search, recency, and forgetting, and do not attempt to explain it further.

Apart from exploration, our analysis also revealed that recency, the behaviour of discounting early experiences, also had a large impact on performance. It predicted that by eliminating recency participants would increase their mean response by 13%. Together, the predicted high impact of exploration and recency on mean response suggests that participants were unsure about how outcomes were generated and tried to learn more about them. Exploration points to this drive to learn more about the environment, and recency indicates that participants believed the environment was non-stationary, which may have resulted from their failing to find a consistent pattern.

### Limitations of the Study

The main limitation of this study is that none of the investigated processes (pattern search, exploration, recency, and forgetting) was manipulated experimentally. All conclusions were drawn by fitting computational models to the data and running simulations based on the obtained results. We also did not attempt to investigate why participants explore, but simply modelled exploratory choices as happening “at random.”

Lastly, it is usually not possible to fit the MPL model to a single participant’s results and obtain precise estimates of the model’s parameters. Our analysis shows that if the *θ* parameter has a low value and an individual fit is attempted, the resulting estimates for the parameters *A*, *ρ*, and *k* are vague. Because of this limitation, it is only possible to work with hierarchical models, which are able to analyse an entire sample of participants at once and obtain much more precise estimates of relevant quantities.

## Conclusion

Our work has thus made novel quantitative and conceptual contributions to the study of human decision making. It confirmed that in a probability learning task the vast majority of participants search for patterns in the outcome sequence, and made the novel estimation that participants believe that each outcome depends on one or two previous ones. But our analysis also indicated that pattern search was not the main cause of suboptimal behaviour: recency and especially exploration had a larger impact on performance. We conclude that suboptimal behaviour in a probability learning task is ultimately caused by participants being unsure of how outcomes are generated, possibly because they cannot find a strategy that results in perfect accuracy. This uncertainty drives them to search for patterns, assume that their environment is changing, and explore.

## Acknowledgements

This work was supported by the São Paulo Research Foundation – FAPESP [grant numbers 2013/10694-0, 2013/13352-2]; the National Council of Technological and Scientific Development – CNPq [grant numbers 132659/2010-7, 305703/2012-9, 248996/2013-4]; and the CAPES Foundation [grant numbers 1587/13-7, 2034/15-8]. Our funding sources had no involvement in study design, in the collection, analysis, and interpretation of data, in the writing of the article, or in the decision to submit it for publication.

MVC Baldo is indebted to the late Prof. Glyn Humphreys for hosting him during a sabbatical at the Oxford Cognitive Neuropsychology Centre and encouraging this work.

## Author contributions statement

CFS and CGV performed the experiments. CFS analysed the data and wrote the computer code, with input from NC and MVCB. CFS, NC, and MVCB developed the computational model. CFS wrote the manuscript and prepared the figures. CFS and MVCB reviewed the manuscript.

## Additional information

The authors declare no competing financial interests.

